# Impairment of a distinct cancer-associated fibroblast population limits tumour growth and metastasis

**DOI:** 10.1101/2020.05.17.100412

**Authors:** Ute Jungwirth, Antoinette van Weverwijk, Liam Jenkins, John Alexander, David Vicente, Qiong Gao, Syed Haider, Marjan Iravani, Clare M. Isacke

**Affiliations:** The Breast Cancer Now Toby Robins Research Centre, The Institute of Cancer Research, London SW3 6JB, UK; Department of Pharmacy and Pharmacology, Centre for Therapeutic Innovation, University of Bath, Bath BA2 7AY, UK; Division of Tumor Biology & Immunology, The Netherlands Cancer Institute, Plesmanlaan 121, 1066 CX Amsterdam, The Netherlands; Strategic Initiatives, Data Science, The Institute of Cancer Research, Sutton SM2 5NG, UK

## Abstract

Profiling studies have revealed considerable phenotypic heterogeneity in cancer-associated fibroblasts (CAFs) present within the tumour microenvironment, however, functional characterisation of different CAF subsets is hampered by the lack of specific markers defining these populations. Here we show that genetic deletion of the Endo180 (*MRC2*) receptor, predominantly expressed by a population of matrix-remodelling CAFs, profoundly limits tumour growth and metastasis; effects that can be recapitulated in 3D co-culture assays. This impairment results from a CAF-intrinsic contractility defect and reduced CAF viability which, coupled with the lack of phenotype in the normal mouse, demonstrates that upregulated Endo180 expression by a specific, potentially targetable CAF subset is required to generate a supportive tumour microenvironment. Further, characterisation of a tumour subline selected via serial *in vivo* passage for its ability to overcome these stromal defects provides important insight into how tumour cells adapt to a non-activated stroma in the early stages of metastatic colonisation.

---

There is extensive functional evidence implicating cancer-associated fibroblasts (CAFs) in tumour progression, via their ability to deposit and remodel the extracellular matrix (ECM), to secrete pro-tumourigenic factors and by modulating the immune compartment^1–5^. However, there is also evidence that stromal fibroblasts can play a role in restraining tumour growth, for example by acting as a desmoplastic barrier to tumour cell invasion and by the recruitment of anti-tumour immune cells^6^. Consideration of how to limit the pro-tumourigenic functions of CAFs, whilst retaining their anti-tumourigenic role, has been hampered by a lack of understanding of the heterogeneity of CAFs within solid tumours and the paucity of specific markers to identify and functionally analyse different CAF subpopulations. Progress in addressing these shortcomings has come from a number of studies reporting single cell sequencing of CAFs or analysis of CAF subsets which has revealed the diversity of fibroblast populations and provided important clues as to their origin and functional properties^7–14^.

In this study, we explore the functional role of a CAF receptor, Endo180 (also known as uPARAP) encoded by the *MRC2* gene. The 180 kDa Endo180 receptor comprises an N-terminal cysteine-rich domain, a fibronectin type II (FNII) domain that has been shown to bind collagens^15–19^, 8 C-type lectin-like domains (CTLDs), of which only CTLD2 displays Ca^2+^-dependent binding of sugars^20^, a single pass transmembrane domain and a cytoplasmic domain that interacts with components of the clathrin-mediated internalisation machinery to drive constitutive recycling between the plasma membrane and intracellular endosomes^21, 22^. Although Endo180 is expressed by some sarcomas^23^, glioblastomas^24^ and metaplastic breast cancers^25^, in most solid tumours of epithelial origin, expression of Endo180 is predominantly restricted to CAFs with little or no expression by the tumour cells^26–29^. Functionally, Endo180 can mediate endosomal uptake of collagens for lysosomal degradation^15, 18^ and has been demonstrated to promote cell migration^16, 17^ via its ability to generate Rho-ROCK-MLC2-mediated contractility^30^.

Here we report that Endo180 expression is associated with the subset of CAFs characterised by their expression of matrix components and matrix-modifying enzymes and that genetic deletion of Endo180, although having little phenotype in normal mice, severely limits the growth and metastatic colonisation of inoculated wildtype syngeneic tumour cells. Mechanistic studies reveal that failure to upregulate Endo180 expression in the tumour microenvironment results in CAFs with impaired contractility and reduced viability, and that this in turn can drive an altered pattern of tumour evolution resulting in the selection of tumour cells with enhanced intrinsic matrix remodelling properties.

## Results

### Expression of Endo180 by cancer-associated fibroblasts (CAFs)

Consistent with the published literature^26–29^ and immunohistochemical data showing that the Endo180 protein is predominantly localised to stromal fibroblasts in solid tumours (Fig. 1a; Supplementary Fig. 1a-c), Endo180 (*MRC2*) expression is restricted to the fibroblast population in tumours separated into EpCAM+ epithelial/tumour cells, CD45+ leukocytes, CD31+ endothelial cells and FAP+ CAFs^31^ (Fig. 1b). In this dataset, Endo180 represents one of the 30 genes whose expression is upregulated in CAFs versus all other populations (Supplementary Fig. 1b). Moreover, when comparing human breast cancers with adjacent normal tissues in the TCGA dataset, there is a trend for increased expression in the tumour samples (Fig. 1c). This increased expression in CAFs does not simply reflect a proportionate increase in the number of fibroblasts in the tumour samples as expression of other classic fibroblast makers, alpha-smooth muscle actin (*ACTA2*) and vimentin (*VIM*), is significantly lower in the tumour samples compared to adjacent normal tissue (Fig. 1c). Furthermore, these data are corroborated by single cell RNA-Seq analysis of stromal cells isolated from mouse melanoma primary tumours and draining lymph nodes (bioRxiv doi.org/10.1101/467225), in which *Mrc2* expression is exclusively found in CAFs but not lymph node fibroblasts or other cell types (Supplementary Fig. 1c). *In vitro* co-culture experiments of MRC5 normal human fibroblasts with human breast cancer cells or mouse mammary cancer cells validated this upregulated expression of Endo180 in activated fibroblasts (Fig. 1d, left panel), with equivalent results seen in published datasets profiling Wi38 or HFF1 human fibroblasts co-cultured with MDA-MB-231 cells^32^ (Fig. 1d, right panels). While there were subtle differences in Endo180 gene expression in breast cancers from the TCGA dataset, divided into the basal-like, HER2-enriched, luminal A and luminal B intrinsic subtypes (1f), fibroblasts isolated from oestrogen receptor positive (ER+), human epidermal growth factor receptor 2-positive (HER2 or triple negative breast cancers (TNBC; lacking expression of ER, progesterone receptor and HER2) showed no difference in Endo180 mRNA levels (Fig.1e) indicating that Endo180 expression in fibroblasts is not breast cancer subtype restricted. Consistent with these data, when taking all breast cancer subtypes together (Fig. 1g) or the intrinsic breast cancer subtypes individually (Supplementary Fig. 1d), there is a strong correlation of Endo180 expression with a fibroblast TGFβ response signature (F-TBRS), that is associated with fibroblast activation in the tumour stroma and is prognostic of poor outcome in colorectal cancers^31, 33^.

**Fig 1.**
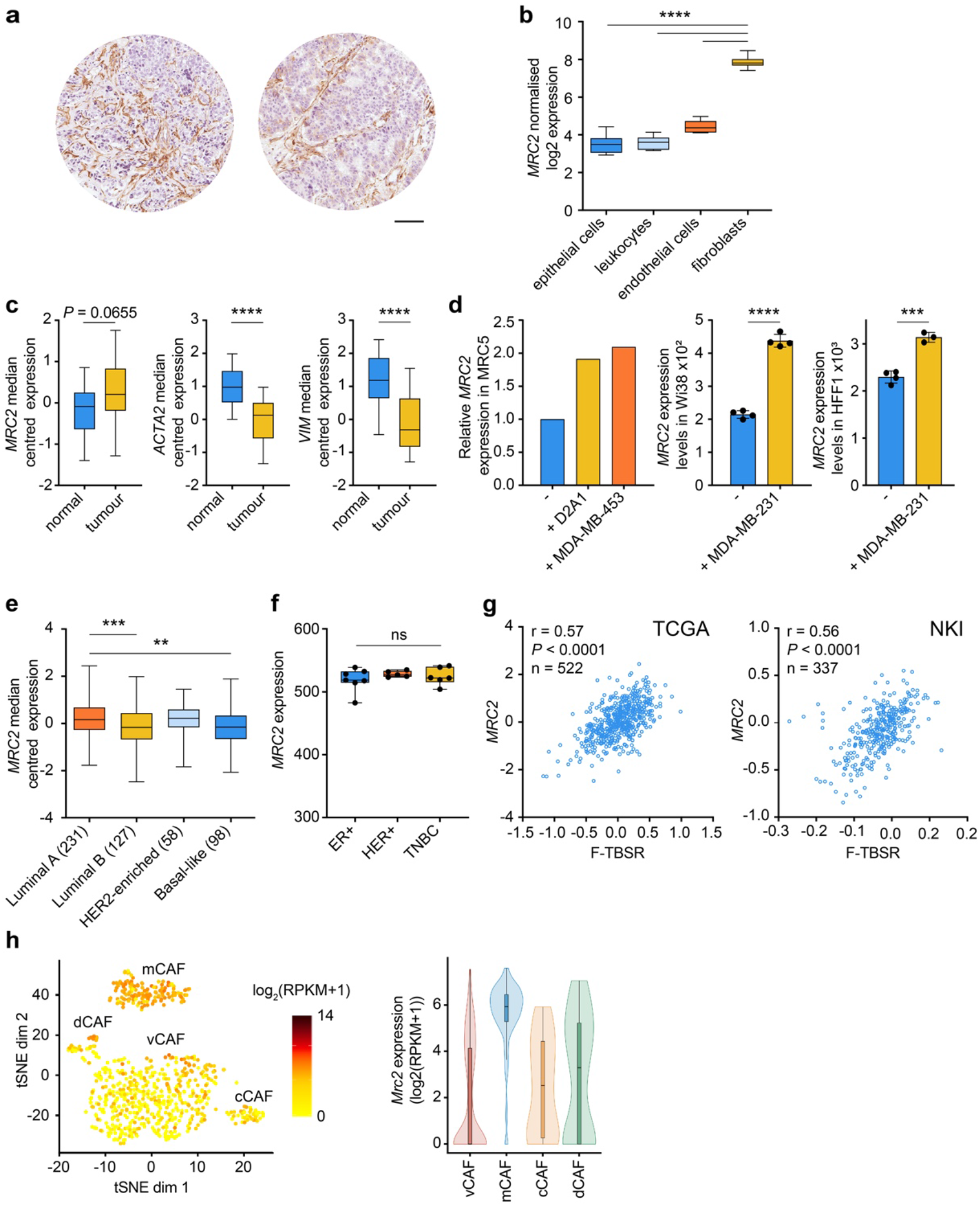
Endo180 is associated with fibroblast activation. **a** Immunohistochemical staining of Endo180 in invasive human breast cancers (scale bar, 0.1 μm). **b** Endo180 (*MRC2*) expression in epithelial cells, leucocytes, endothelial cells and fibroblasts purified from human colorectal cancers (GSE39397^31^. Tumours from 6 individual patients (one-way ANOVA with Tukey’s multiple comparison test). **c** Boxplots of Endo180 (*MRC2)*, *ACTA2* and *VIM* expression in the TCGA dataset of adjacent normal and breast cancer tissue (n=22 per group; paired *t*-test). **d** Left panel, Relative Endo180 (*MRC2*) expression (*B2M* endogenous control) in human MRC5 fibroblasts cultured alone or co-cultured with D2A1 or MDA-MB-453 tumour cells. Middle and right panels, Endo180 (*MRC2*) expression in Wi38 and HFF1 human fibroblasts, respectively, cultured alone or co-cultured with MDA-MB-231 (GSE41678^32^; n=4 per group; *t*-test). **e** Endo180 (*MRC2*) expression in TCGA breast cancer intrinsic subtypes (one-way ANOVA with Tukey’s multiple comparison test). Numbers of samples in each category are indicated. **f** Endo180 (*MRC2*) expression in fibroblasts isolated from different breast cancer subtypes (GSE37614^66^; n=5-7 per group; one-way ANOVA with Tukey’s multiple comparison test). **g** Correlation of Endo180 (*MRC2*) gene expression with a fibroblast TGFβ response signature (F-TBRS)^31^ in breast cancers from the TCGA (Pearson correlation, r=0.57, *P* < 0.0001 n=522) and the NKI295^67^ (Pearson correlation, r=0.56, *P* < 0.0001, n=337) datasets. **h** Endo180 (*Mrc2*) gene expression in the single cell sequencing dataset of CAFs isolated from MMTV-PyMT tumours^7^. Shown is *Mrc2* expression log_2_(RPKM+1) in individual cells (left panel, tSNE plot) and in the vascular (vCAF), matrix (mCAF), cycling (cCAF) and development (dCAF) subpopulations (right panel, violin plots).

As well as inter-tumour heterogeneity within the stromal compartments of solid cancers, there is increasing evidence of intratumoural stromal heterogeneity, including CAF heterogeneity^7–14^. For example, single cell sequencing of fibroblasts isolated from the mammary tumours in MMTV-PyMT transgenic mice ^7^ identified 3 fibroblast subsets defined as vascular CAFs, matrix CAFs and developmental CAFs plus a group of actively cycling vascular CAFs termed cycling CAFs. Endo180 is most strongly expressed by matrix CAFs (Fig. 1h), the subset characterised by expression of ECM and ECM-related genes and predicted to originate from activated resident fibroblasts. These data indicate that Endo180 expression is upregulated in a discrete population of CAFs in solid tumours.

### Endo180 expression is required for efficient tumour growth and metastasis

To investigate the functional consequence of the upregulated Endo180 expression by matrix CAFs, mice with a targeted deletion in Endo180 (*Mrc2*)^17^ were backcrossed onto a BALB/c background (see Methods) and inoculated orthotopically with syngeneic 4T1 mouse mammary cancer cells. In the wildtype mice, 4T1 tumours grew at the expected rate reaching maximal allowable size on day 30, and examination of the lungs post-mortem revealed readily detectable spontaneous macrometastatic disease in 5 out of 6 mice. By contrast, primary tumours only developed in 3 out of 6 Endo180^−/−^ mice and those mice that did develop tumours showed a delay in tumour growth and a significant reduction in spontaneous metastasis (Fig. 2a). To confirm these data, the experiment was repeated with the D2A1 mouse mammary tumour cell line (n= 9 mice per genotype). Whereas, primary tumours readily grew in 8 of the wildtype mice, tumour take was only observed in 4 of the Endo180^−/−^ mice and, again, the tumours that did develop in the Endo180^−/−^ mice grew more slowly such that at the termination of the experiment, there was a significant difference not only in tumour volume but also tumour weight. D2A1 cells are poorly metastatic and therefore an impairment in metastasis could not be ascertained, however, we recently derived the D2A1-m2 subline by selecting for enhanced metastatic growth via serial *in vivo* passage in wildtype BALB/c mice^34^. After orthotopic inoculation, the D2A1-m2 primary tumours grew equally well in both wildtype and Endo180^−/−^ mice but there remained a significant defect in their ability to spontaneously metastasise in the Endo180^−/−^ mice (Fig. 2c, lower panels), confirming that all three cell lines examined show impaired tumour progression but that the defects manifested vary between the lines. Moreover, the defect in primary tumour take and growth in the 4T1 and D2A1 lines and the defect in spontaneous metastasis in the more aggressive 4T1 and D2A1-m2 lines suggested an inability of Endo180 to activate their tumour stroma to support the initial stages of tumour growth.

**Fig 2.**
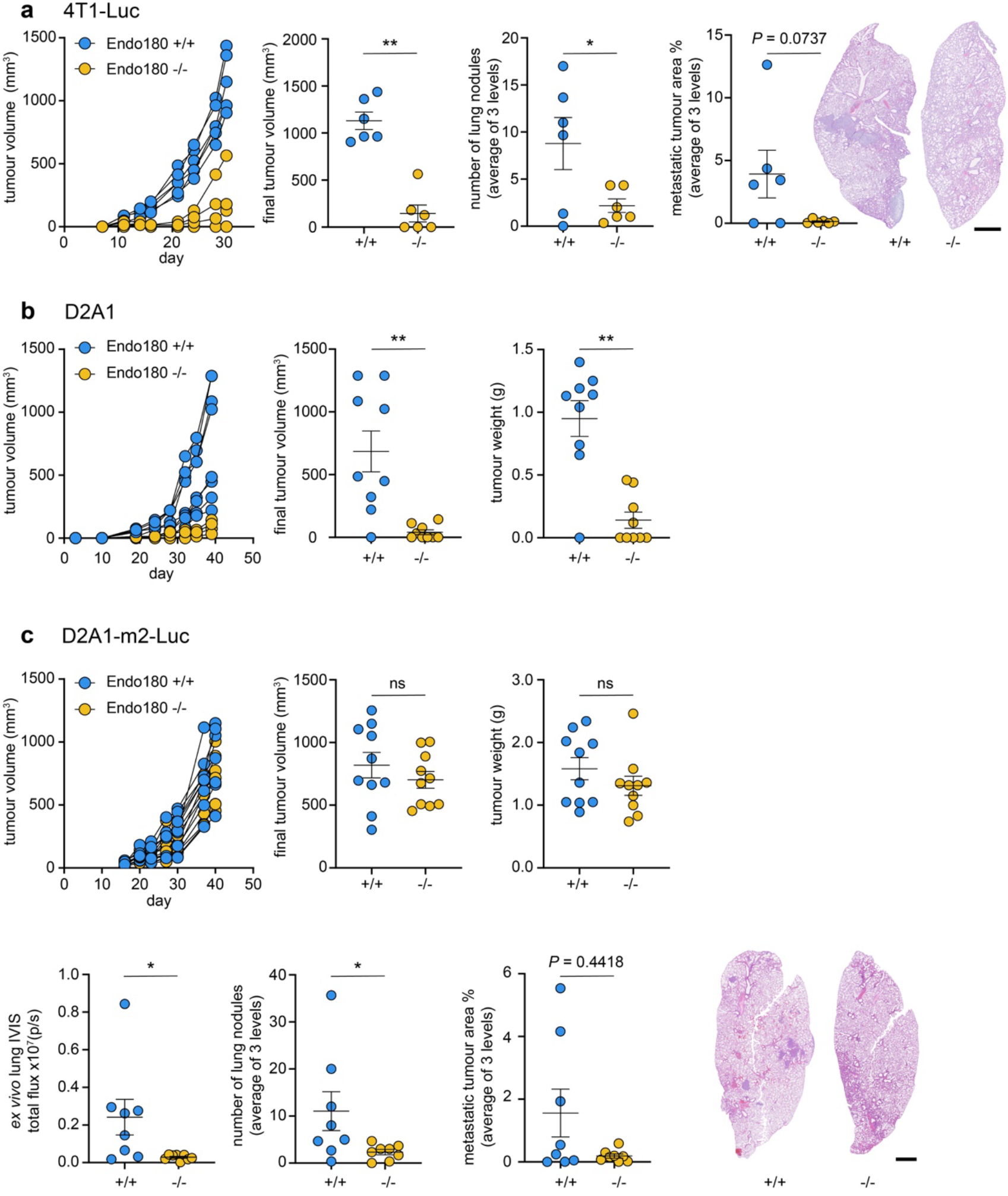
Endo180 promotes tumour take, growth and spontaneous metastasis. Tumour cells were injected orthotopically into the 4^th^ mammary fat pad of either BALB/c Endo180^+/+^ or Endo180^−/−^ mice. Tumour volume was measured twice weekly. Spontaneous lung metastasis at the end of the experiment was quantified as mean number of lung nodules and mean % of tumour area in 3 lung sections. All data shown represents mean values ±SEM. Representative lung sections (scale bar, 1 mm). **a** 1 × 10^4^ 4T1-Luc-RFP cells inoculated (n=6 mice per group). Shown are; primary tumour growth in individual mice (no tumours were detected in 3 Endo180^−/−^ mice), final tumour volume at day 30 (Mann-Whitney *U* test), quantification of spontaneous metastasis lung metastatic burden by number of lung nodules (*t*-test) and % metastatic area (*t*-test) and representative H&E images. **b** 5 × 10^4^ D2A1 cells inoculated (n=9 mice per group). Shown are primary tumour growth in individual mice (no tumours were detected in 4 out of 9 Endo180^−/−^ and 1 out of 9 Endo180^+/+^ mice), final tumour volume at day 39 and tumour weight (both Mann-Whitney *U* test). **c** 5 × 10^4^ D2A1-m2-Luc cells inoculated (n=10 mice per group). Shown are; primary tumour growth, final tumour volume at day 40 (*t*-test), tumour weight (*t*-test), mice with intraperitoneal tumour growth were excluded in subsequent metastasis analysis (2 in each group), representative H&E images of the lungs, quantification of spontaneous metastasis to the lung by *ex vivo* IVIS imaging (Mann-Whitney *U* test), number of lung nodules (*t*-test) and % metastatic area (Mann-Whitney *U* test).

To address this directly, tumour cells were inoculated via the tail vein into recipient mice, which results predominantly in single cells seeding into the lungs. BALB/c mice inoculated with 4T1 cells (Fig. 3a; Supplementary Fig. 2a), the related, but less aggressive, 4T07 cells (Fig. 3b; Supplementary Fig. 2b), D2A1 cells (Fig. 3c; Supplementary Fig. 2c) and D2A1-m2 cells (Fig. 3d) all developed large macrometastatic lesions in the lungs of wildtype mice but were severely impaired in their growth in Endo180^−/−^ mice as monitored by IVIS imaging for luciferase tagged cells, *ex vivo* lung weight or quantification of tumour burden in histological sections. Importantly, equivalent results were obtained following intravenous inoculation of C57BL/6 Endo180^−/−^ mice with the syngeneic E0771 mouse mammary carcinoma line (Fig. 3e; Supplementary Fig. 2d) demonstrating that the defect in lung colonisation is not restricted to the BALB/c background. Finally, we also investigated whether this effect was evident in other organs by performing intrasplenic inoculation of 4T1 cells.

**Fig 3.**
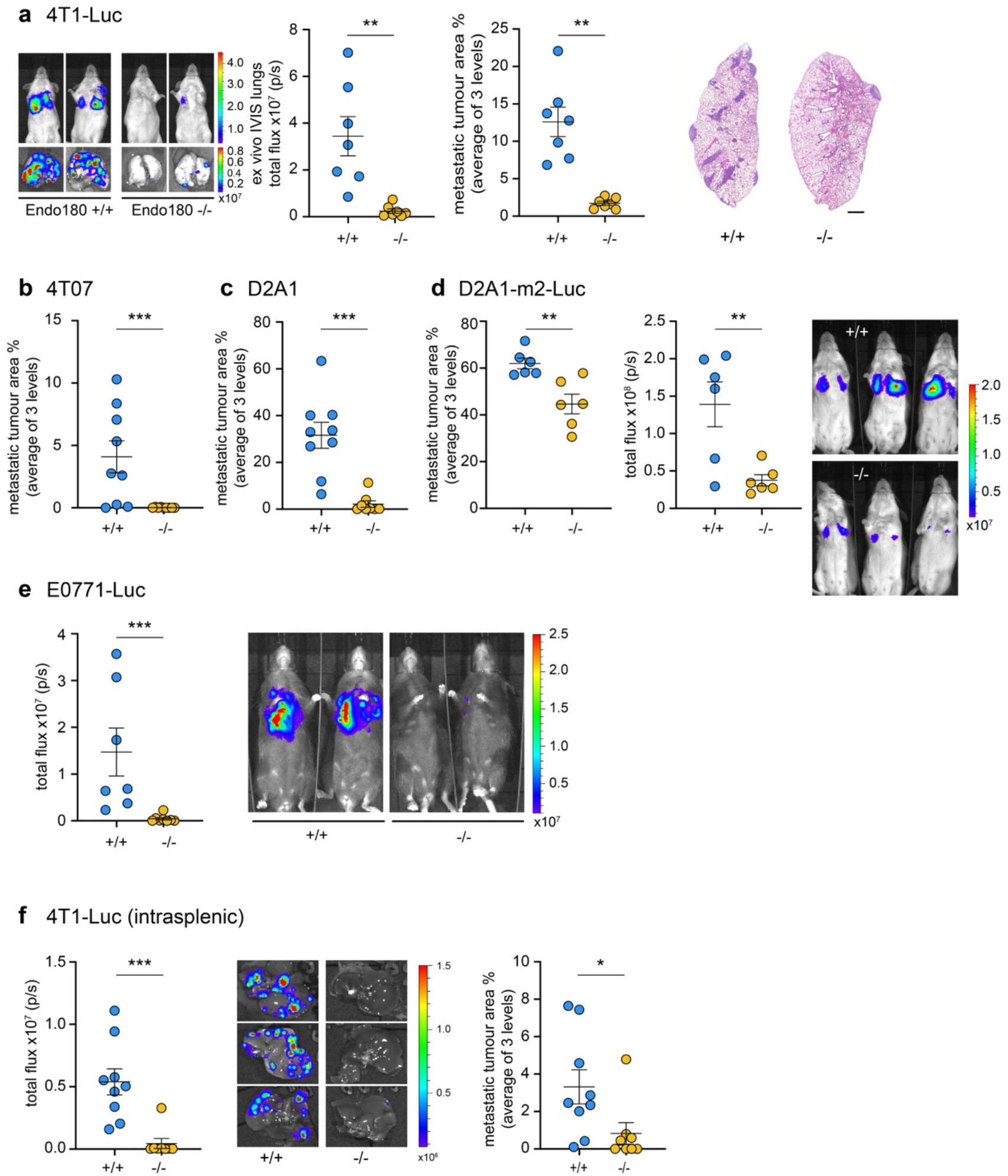
Endo180 promotes metastatic tumour growth in lung and liver. Additional quantification and images are provided in Supplementary Fig. 2. **a** 1 × 10^5^ 4T1-Luc cells injected intravenously into BALB/c mice (n=7 per group). The experiment was terminated on day 12. Shown are; representative *in vivo* and *ex vivo* IVIS imaging, quantification of *ex vivo* lung IVIS imaging (*t*-test), quantification of metastatic tumour area (*t*-test) and representative H&E stained lung sections. **b** 2.5 × 10^5^ 4T07 cells injected intravenously into BALB/c mice (n=9 per group). The experiment was terminated on day 24. Shown is quantification of metastatic tumour area in the lungs (Mann-Whitney *U* test). **c** 4 × 10^5^ D2A1 cells injected intravenously into BALB/c mice (n=8 per group). The experiment was terminated on day 23. Shown is quantification of metastatic tumour area in the lungs (Mann-Whitney *U* test). **d** 4 × 10^5^ D2A1-m2-Luc cells injected intravenously into BALB/c mice (n=6 per group). The experiment was terminated on day 14. Shown are representative *in vivo* IVIS images, quantification of *ex vivo* lung IVIS imaging (*t*-test) and quantification of metastatic tumour area in the lungs (*t*-test). **e** 4 × 10^5^ E0771-Luc cells injected intravenously into C57BL/6 mice (n=9 or 10 per group). The experiment was terminated on day 17. Shown are representative *in vivo* IVIS images, and quantification of *in vivo* lung IVIS imaging (Mann-Whitney *U* test). **f** 1 × 10^5^ 4T1-Luc cells injected into the spleen parenchyma of BALB/c mice (n=8 or 9 per group). The experiment terminated on day 13. Shown are; representative *ex vivo* IVIS images of the liver, quantification of *ex vivo* liver IVIS imaging (*t*-test) and quantification of tumour area in the liver (Mann-Whitney *U* test). All data are mean values ±SEM. IVIS data are mean total flux values from organs imaged *ex vivo*. Metastatic burden was quantified as mean % of tumour area in 3 lung or liver sections.

Whereas all wildtype mice developed liver metastases, only 1 out of 8 Endo180^−/−^ mice showed evidence of macrometastatic disease (Fig. 3f; Supplementary Fig. 2e).

Together these data provide strong evidence that upregulated expression of Endo180 in matrix CAFs is required for efficient tumour progression *in vivo*. If this assertion is correct, then there should be no difference in tumour colonisation in tissues that lack stromal fibroblasts. Endo180 expression is not detected in the normal brain apart from weak expression in some perivascular cells ^24^ and similarly there is lack of expression of classic fibroblast markers such as FSP1 and αSMA (Supplementary Fig. 3a). Interestingly, following direct intracranial inoculation of 4T1 tumours cells there was no significant difference in brain tumour burden in wildtype and Endo180^−/−^ mice (Supplementary Fig. 3b). Similarly, there was no significant difference in brain metastasis following intracardiac inoculation of 4T1 cells (Supplementary Fig. 3c).

### *In vitro* assays recapitulate *in vivo* phenotypes

To investigate the mechanism by which Endo180 promotes tumour progression, fibroblasts in which Endo180 expression had been downregulated by siRNA transfection or transduction with shRNA constructs (Supplementary Fig. 4) were co-seeded with tumour cells into low adherence U-bottom plates resulting in the formation of 3D coculture spheroids. Growth, as monitored by tumour area and CellTiter-Glo viability, of D2A1 tumour cell spheroids was significantly enhanced when admixed with mouse CAFs, immortalised normal mouse mammary gland fibroblasts (NF#1) or NIH-3T3 fibroblasts (Fig. 4a). By contrast, admixing with fibroblasts that had been transfected with siRNAs targeting Endo180 severely impaired fibroblast-mediated enhanced spheroid growth. Equivalent results were obtained with 4T1 mouse mammary carcinoma cells, D2A1 cells admixed with CAFs at different rations (Supplementary Fig. 5a) and with D2A1 cells admixed with mouse CAFs transduced with two independent shRNAs targeting Endo180 (Fig. 4b; Supplementary Fig. 5b,c).

**Fig 4.**
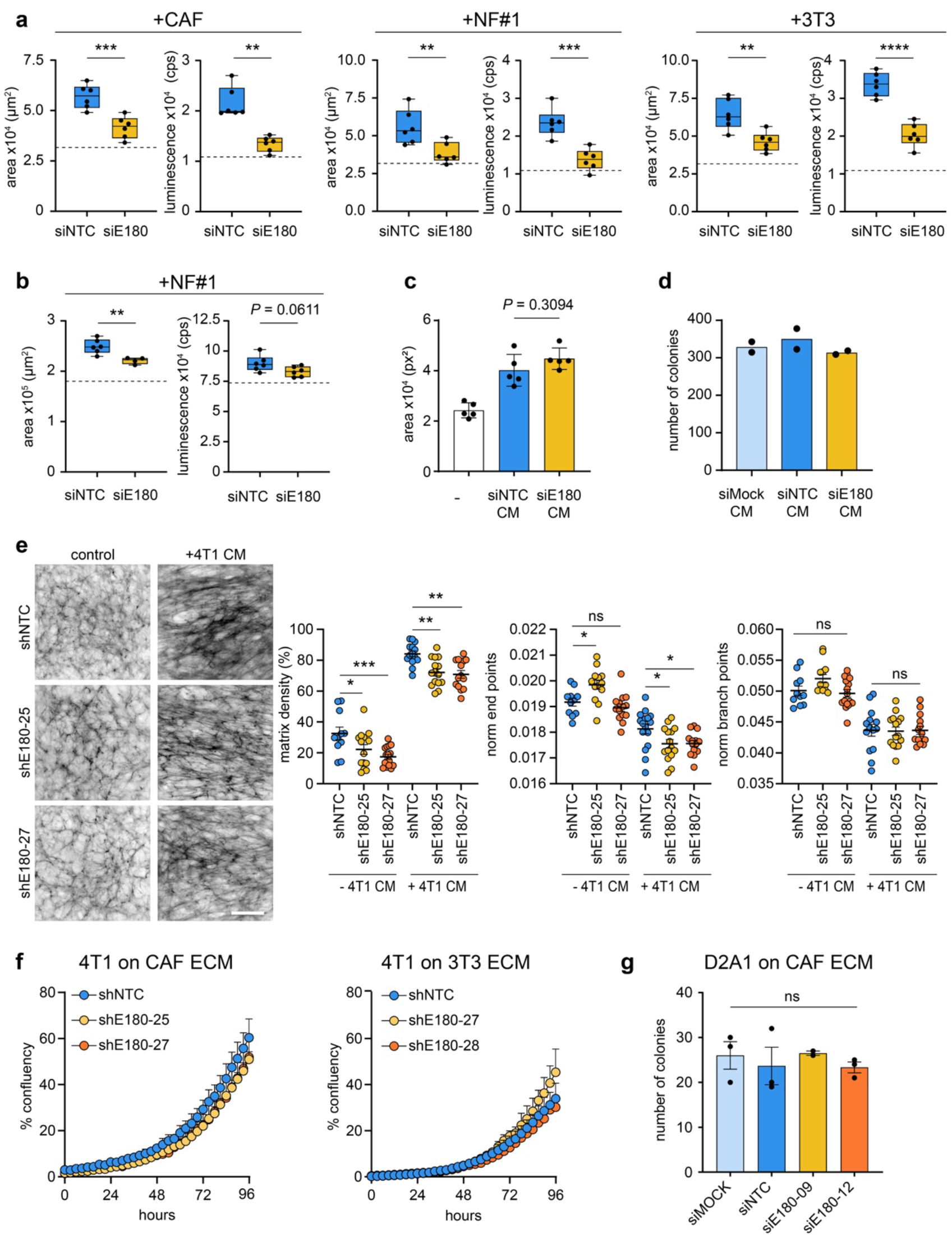
Spheroid co-cultures recapitulate *in vivo* Endo180 impact. **a,b** Fibroblast-tumour cell co-culture spheroid assay. **a** 300 D2A1 cells alone or in combination with 600 mouse CAFs, NF#1 or 3T3 fibroblasts transfected with non-targeting control (NTC) or Endo180 (E180) targeting siRNA oligonucleotides were seeded into U-bottom low adherence plates and cultured for 8 days. Spheroid growth was assessed by spheroid area and CellTiter-Glo. Box plot shows median, 25^th^ to 75^th^ quartile, whiskers show minimum and maximum, dotted line shows mean spheroid size/CellTiter-Glo value of tumour cells alone (n=6 spheroids per condition; mean values ±SEM, *t*-test). **b** 4T1 spheroid growth assessed by spheroid area and CellTiter-Glo when cultured alone (dotted line) or co-cultured with siNTC or siEnd180 NF#1 as described in panel a (n=6 spheroids per condition; mean values ±SEM, *t*-test). **c** D2A1 spheroid growth in DMEM plus 2% FBS or serum-free conditioned medium (CM) collected from CAFs transfected with siNTC or siEndo180 and then supplemented with 2% FBS (n=3-6 spheroids; mean values ±SEM, one-way ANOVA, Tukey post-hoc test). **d** D2A1 colony formation assay in the presence of CM collected from Mock, siNTC or siEndo180 transfected CAFs and supplemented with 2% FBS (n=2). **e** CAFs transduced with shNTC or two independent shRNAs targeting Endo180 cultured with or without CM from 4T1 cells supplemented with 2.5% FBS. Cultures were stained for fibronectin. Left panel, representative images of fibronectin fibres (scale bar, 100 μm). Right panels, quantification of matrix density, normalised endpoints and normalised branch points (n=16 fields of view per condition; mean values ±SEM, two-way ANOVA, Tukey post-hoc test). **f** 4T1 cell growth, measured by confluency, on ECM derived from CAFs or 3T3 fibroblasts transduced with shNTC or two independent shRNAs targeting Endo180 (n=3 wells per condition; mean values ±SEM, two-way ANOVA, non-significant in all comparisons). **g** D2A1 colony formation assay on ECM generated from mock transfected CAFs or CAFs transfected with siNTC or two independent siRNAs targeting Endo180 (n=3; mean values ±SEM, one-way ANOVA).

**Fig 5.**
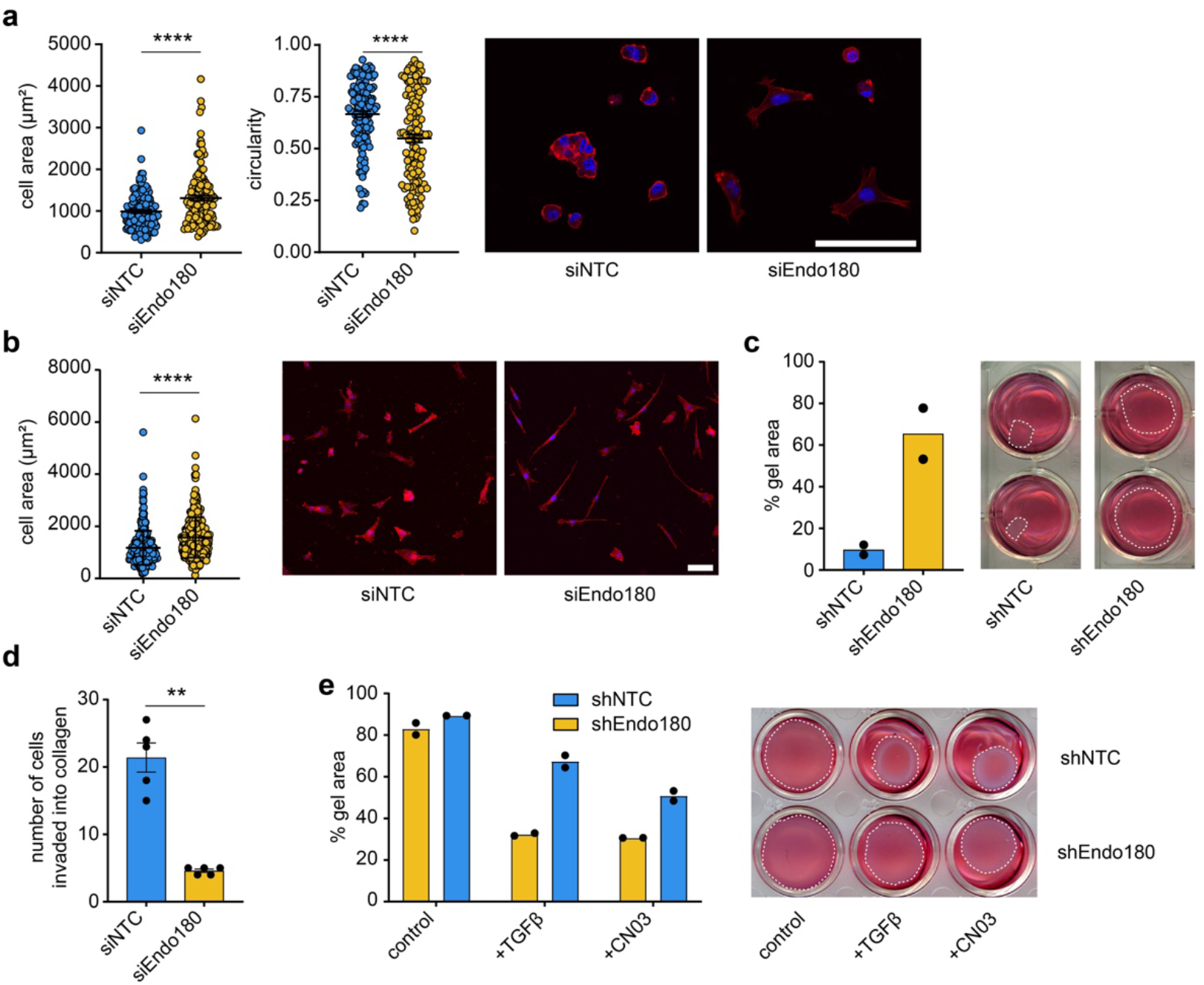
Loss of contractility and hypersensitivity. **a,b** Cell spreading (area) and cell circularity of siNTC and siEndo180 CAFs cultured on soft 2 kPa hydrogels coated with (**a**) collagen (n=134 cells; mean values ±SEM, Mann-Whitney *U*) or (**b**) fibronectin (n=300 cells; mean values ±SEM, Mann-Whitney *U* test). Cells were stained with DAPI and Alexa555-phalloidin. Right panels show representative images (scale bar, 100 μm). **c** shNTC or shEndo180 transfected CAFs were embedded into rat tail collagen (2 mg mL^−1^). Data show % gel area (n=2) after 9 days. Right panel, representative images. **d** Number of siNTC or siEndo180 CAFs invading from fibroblast spheroids into a collagen matrix (2 mg mL^−1^) after 24 hours (n=5; mean values ±SEM, *t*-test). **e** shNTC or shEndo180 3T3 fibroblasts were embedded in collagen gels and treated with vehicle, TGFβ and/or CN03. Data show % gel contraction after 24 hours (n=2).

These data indicate that failure to express Endo180 severely impairs the ability of fibroblasts to promote tumour cell proliferation in 3D spheroid culture.

Interestingly, conditioned medium from wildtype CAFs promoted the growth D2A1 tumour spheroids, however, conditioned medium from siEndo180 transfected CAFs was equally as effective (Fig. 4c), indicating that Endo180 expression does not regulate expression or release of soluble tumour growth promoting components. Similarly, there was no difference in D2A1 2D colony formation in the presence of mock, siNTC and siEndo180-transfected CAF conditioned medium (Fig. 4d). As Endo180 has been described as a matrix remodelling receptor based on its ability to internalise collagens for intracellular degradation^15, 18^ and is expressed predominantly by matrix CAFs (Fig. 1h) we next addressed whether the ECM produced by Endo180 wildtype and Endo180-deficient CAFs differentially modulated tumour cell growth. CAFs transduced with a non-targeting shRNA or two independent shRNAs targeting Endo180 were cultured alone or with conditioned medium from 4T1 tumour cells to generate CAF-derived ECM. Staining of the decelluarised matrices for fibronectin allowed an examination of the ECM deposition and fibre alignment using the TWOMBLI macro in FIJI (bioRxiv doi.org/10.1101/867507). Endo180-deficient cells cultured alone or with 4T1 conditioned medium produced a reduced density matrix (Fig. 4e, left panels). As well as resulting in the production of a denser matrix, treatment of both wildtype and Endo180-depeleted CAFs with 4T1-conditioned medium resulted in a more anisotropic matrix, indicated by the reduced number of normalised end and branching points (Fig. 4e right panels). However, there was no difference in anisotropy in Endo180-deficient compared to the Endo180 wildtype CAF-derived matrices and while tumour cells grew better on a CAF-derived ECM compared to a 3T3 fibroblast-derived ECM (Supplementary Fig. 5c), there was again no difference in tumour cell proliferation (Fig. 4f, Supplementary Fig. 5c) or tumour cell colony formation number (Fig. 4g) on matrices derived from Endo180 wildtype or Endo180-deficient CAFs.

### Endo180-deficient fibroblasts show reduced viability in the activated tumour stroma

As neither fibroblast conditioned medium nor fibroblast-derived ECM could recapitulate the inability of Endo180-deficient fibroblasts to support efficient tumour cell growth, we next addressed whether this defect was intrinsic to the fibroblasts. When plated onto collagen-coated soft (2 kPa) hydrogels, wildtype CAFs were able to contract their cytoskeleton and round up, whereas Endo180-deficient cells remained spread resulting in a significantly increased single cell area and decreased circularity (Fig. 5a). This effect was not specific to a collagen substratum as equivalent results were observed with cells plated onto fibronectin-coated hydrogels (Fig. 5b). Equivalent results were obtained with NF#1 and 3T3 fibroblasts and with stiff hydrogels (Supplementary Fig. 6a,b). Consistent with this observed contractile defect, when embedded into a collagen matrix, Endo180-deficient CAFs show a striking inability to efficiently contract the collagen gel (Fig. 5c) or to invade into a 3D matrix (Fig. 5d; Supplementary Fig. 6c). To determine whether the contractility defect of Endo180-deficient fibroblasts could be overcome, fibroblasts in 3D culture were treated with TGFβ, a key driver of fibroblast activation both *in vitro* and *in vivo*^31^. When plated into collagen matrices, TGFβ pre-treatment dramatically promoted collagen gel contraction by wildtype 3T3 fibroblasts but had little impact on the ability of Endo180-deficient fibroblasts to contract the gels (Fig. 5e). Similarly, treatment with the cell permeable Rho activator CN03^35, 36^ enhanced the collagen gel contractility of wildtype, but not Endo180-deficient, fibroblasts, consistent with the ability of internalised Endo180 to promote localised Rho-ROCK-MLC2 contractile signals^30^.

**Fig 6.**
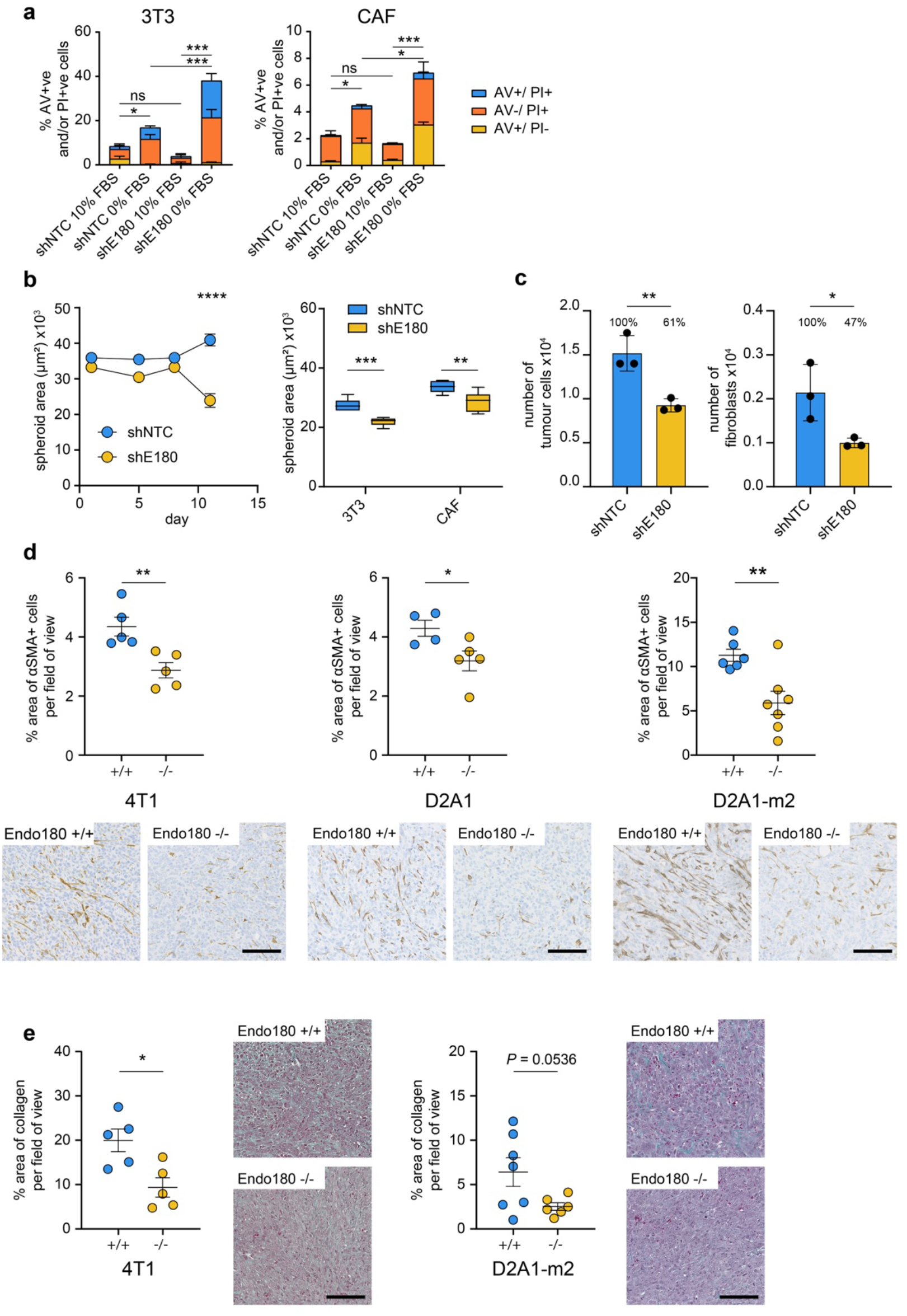
Reduced fibroblast viability. **a** Cell survival under anoikis conditions (flat bottom low adherence 6-well plates) in 0 or 10% FBS measured by Annexin V (AV)/ PI staining. Left panel, shNTC and shEndo180 3T3, Right panel CAFs. (n=3; mean values ±SEM, one-way ANOVA of total % AV+ve and/or PI+ve cells). **b** Left panel, 900 shNTC or shEndo180 CAF were plated into U-bottom wells and growth of suspension aggregates monitored over time (n=6; mean values ±SEM, two-way ANOVA). Right panel, 900 CAF and 3T3 were plated into U-bottom wells and suspension aggregate size measured on day 10 (n=6; mean values ±SEM, *t*-test). **c** 300 D2A1 and 600 GFP-CAFs cells were co-cultured in U-bottom plates. After 8 days, spheroids were dissociated and the number of GFP-positive CAFs counted (n=3 per condition; mean values ±SEM, *t*-test). **d** Analysis of αSMA positive staining in 4T1, D2A1 and D2A1-m2 primary tumours grown in wildtype and Endo180^−/−^ mice. Data represents mean values per tumour ±SEM (*t*-test) Representative images (scale bar, 100 μm). **e** Analysis of Masson’s trichrome staining of 4T1 (left panel) and D2A1-m2 (right panel) primary tumours in wildtype and Endo180^−/−^ mice. Data represent mean values per tumour ±SEM (*t*-test). Representative images (scale bar, 100 μm).

Together these findings raise the question as to whether the reduced tumour growth and metastasis observed in the Endo180^−/−^ mice is directly linked to an altered biology associated with the contraction-defective Endo180^−/−^ fibroblasts or that a contractile defect has a cell intrinsic effect on fibroblast viability in the tumour stroma. In the support of the latter hypothesis, Endo180-deficient fibroblasts plated into non-adherent conditions in low serum, but not 10% serum, have an increased level of apoptosis (Fig. 6a). Similarly, while wildtype CAFs plated alone in U-bottom plates form stable 3D aggregates that showed a modest increase in size over time, Endo180-deficient CAF 3D aggregates deteriorated such that they were significantly smaller at day 10 (Fig. 6b). As a consequence, we assessed whether the defect in tumour spheroid growth in the presence of Endo180-deficient CAFs resulted from a differential loss of CAFs in the 3D cultures. Dissociation of 3D D2A1 tumour spheroids admixed with CAFs revealed, a significantly lower number of Endo180-deficient CAFs retrieved compared to wildtype CAFs (Fig. 6c). Consistent with these findings, analysis of 4T1, D2A1 and D2A1-m2 primary tumours revealed a significant deficit in αSMA positive cells in the tumour stroma of Endo180^−/−^ recipient mice (Fig. 6d). Moreover, these tumours also displayed a reduction in fibrillar collagen as evidenced by Masson’s trichrome staining suggesting that the reduced CAF numbers in the Endo180^−/−^ mouse tumours result in reduced collagen deposition (Fig. 6e).

#### Tumour adaptability to stromal deficiency

Tumours can evolve to overcome therapeutic insult or the lack of a supportive microenvironment^37^ and consequently it was important to investigate how tumours might adapt to overcome the microenvironmental block associated with loss of Endo180. Previously we have described the selection of metastatic sublines from the non-metastatic D2A1 mouse mammary tumour cell line by repeated *in vivo* passage in wildtype BALB/c mice^34^. Here we used a similar approach to generate a D2A1 subline, D2A1-m12, by serial *in vivo* passage in Endo180-deficient mice (see Methods). Compared to the parental D2A1 cells, the selected D2A1-m12 subline rapidly formed macrometastatic lesions in the Endo180^−/−^ BALB/c mice following intravenous inoculation (Fig. 7a) and, in contrast to the parental and other mouse mammary cell lines tested (Fig. 3), gave rise to an equivalent metastatic burden when inoculated intravenously into Endo180^−/−^ and wildtype mice (Fig. 7b). Moreover, in a spontaneous metastasis experiment D2A1-m12 primary tumours showed an increased growth rate in Endo180^−/−^ mice, albeit with no significant difference tumour weight at necropsy, and no difference in the metastatic burden in the lungs (Supplementary Fig. 7a).

**Fig 7.**
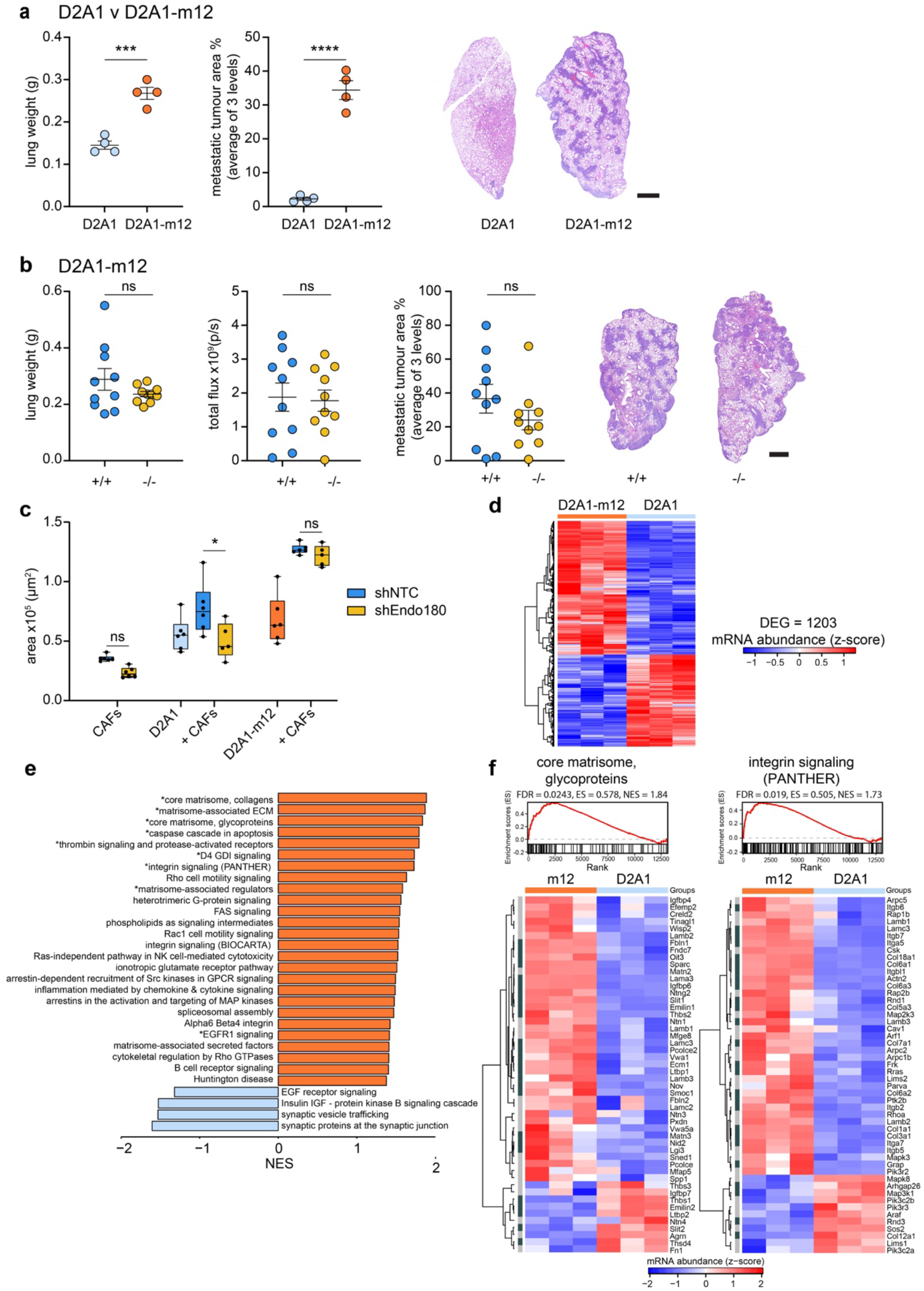
Selected tumour sublines can overcome a defect in stromal Endo180. **a** 4 × 10^5^ D2A1 or D2A1-m12-Luc cells were injected intravenously into BALB/c Endo180^−/−^ mice (n=4 mice per group) and the experiment was terminated on day 12. Shown are *ex vivo* lung weights (*t*-test), quantification of metastatic tumour area in the lungs (*t*-test) and representative H&E stained lung sections (scale bar, 1 mm). **b** 4 × 10^5^ D2A1-m12-Luc cells were injected intravenously into BALB/c Endo180^+/+^ or Endo180^−/−^ mice (n=8 per group) and the experiment terminated on day 12. Shown are *ex vivo* lung weights (*t*-test), *ex vivo* IVIS imaging of the lungs (*t*-test) quantification of metastatic tumour area in the lungs (*t*-test) and representative H&E stained lung sections (scale bar, 1 mm). **c** Tumour cells (D2A1 or D2A1-m12) and/or shNTC/shEndo180 CAFs were cultured in U-bottom plates and spheroid area measured at day 8 (n=6 spheroids per group, 2 way-ANOVA, Sidak post-hoc test). **d** Heatmap displaying mRNA abundance of all 1203 significantly differentially expressed genes (DEGs) determined by RNA-Seq, in D2A1-m12 compared with D2A1 cells (n = 3; heatmap scale is a z-score). Threshold for differential expression was |log_2_FC| > 1, and adjusted *P* value ≤ 0.05. **e** fGSEA, upregulated (orange) and downregulated (blue) in D2A1-m12 compared with D2A1 cells. All pathways displayed had *P* value < 0.05 and those labelled with a * had FDR adjusted P value < 0.1. NES, normalised enrichment score. **f** fGSEA of the pathways ‘core matrisome, glycoproteins’ and ‘integrin signalling pathway (PANTHER)’ and associated heatmap showing the genes of the pathway (n = 3; heatmap scale is a z-score). Dark grey, significant DEGs with |log_2_FC| > 1 and adjusted *P* value ≤ 0.05. Light grey, nonsignificant DEGs.

In *in vitro* assays, admixing D2A1 or D2A1-m12 cells with CAFs significantly promoted 3D co-culture spheroid growth (Fig. 7c). However, as reported in Fig. 4a, whereas Endo180-deficient CAFs were impaired in their ability to promote the growth of D2A1 3D co-culture spheroids there was no significant difference in D2A1-m12 spheroid growth when admixed with wildtype or Endo180-deficient fibroblasts (Fig. 7c). Moreover, whereas D2A1 cells admixed with either wildtype or Endo180-deficient fibroblasts were unable to effectively mediate collagen-gel contraction, gel contraction was much more pronounced with the D2A1-m12 cells even when admixed with Endo180-deficient fibroblasts (Supplementary Fig. 7b). Together these data support the notion that tumour cell selection can generate sublines that have overcome, at least in part, the requirement for fibroblast support both *in vitro* and *in vivo*. To address the mechanism by which this may occur, D2A1 and D2A1-m12 were subject to whole exome sequencing and transcriptional profiling. Exome sequencing revealed that when compared to a reference BALB/c genome, the D2A1 and D2A1-m12 cell lines display a similar copy number aberration profile (Supplementary Fig. 7c) as expected given that the D2A1-m12 subline is derived from the D2A1 parental cells. Comparison of their mutation profiles revealed a similar mutation burden in the D2A1 and D2A1-m12 cell lines (Supplementary Fig. 7d,e). Although there was a significant overlap of shared mutations (Supplementary Fig. 7e), both cell lines also carried distinct mutations, consistent with the parental D2A1 containing a heterogenous mix of subclones and the D2A1-m12 subline being derived from a subset of these during the *in vivo* passage. RNA-Seq analysis of the D2A1 and D2A1-m12 cells revealed 1203 significantly differentially expressed genes (Fig. 7d, Supplementary Fig. 7f), including a 2.96-fold increase in αSMA (*Acta2*) levels. We performed fast Gene Set Enrichment Analysis (fGSEA) using GSKB pathways supplemented with the mouse core matrisome and matrisome-associated pathways from the Matrisome Project due to the lack of well annotated ECM components in most gene ontology categories^38^ (Fig. 7e; Supplementary Table 1). Notably, within the 9 significantly enriched (FDR adjusted *P* value < 0.1) upregulated pathways were core matrisome collagens, core matrisome glycoproteins, matrisome-associated glycoproteins and integrin signalling (Fig. 7e,f; Supplementary Fig. 7g); suggesting that the enhanced tumourigenesis of the D2A1-m12 subline in Endo180^−/−^ mice results from tumour cells acquiring an enhanced ability to remodel and respond to the ECM, when CAF activation and viability is impaired.

## Discussion

CAFs are one of the most abundant cell types within the tumour stroma, however it is now clear from the extensive transcriptional profiling of CAF subsets and, more recently, single cell sequencing studies that the term CAF represents a phenotypically and functionally diverse mix of cells. Traditionally, CAFs have been proposed to promote tumourigenesis via their ability to deposit and remodel the ECM and by their ability to interact both with tumour cells and other stromal cell types via the secretion of growth factors, chemokines and cytokines. However, both normal stromal fibroblasts and CAFs have also been accredited with tumour suppressive properties and it remains unclear how tumour cells orchestrate their interactions with the microenvironment to overcome fibroblast/CAF mediated suppression and take advantage of the pro-tumourigenic CAF functions. One explanation for observed CAF diversity in tumours is that they might have different origins^5, 6, 39, 40^. Certainly, CAFs can arise from the activation of resident stromal fibroblasts, and indeed these may well represent the bulk of the CAFs associated with early lesions. In addition, there is evidence that CAFs can originate from mesenchymal stem cells (MSCs) and, in some mouse models, bone marrow-derived MSCs have been demonstrated to contribute to the tumour stroma. In addition to a bone marrow origin, perivascular cells from a wide variety of tissues have been demonstrated to exhibit phenotypic and functional properties associated with MSCs^41, 42^. In this respect it is of interest that in their CAF single cell sequencing study, despite excluding cells expressing the pericyte marker NG2, Bartoschek and colleagues defined a large population of fibroblasts as ‘vascular CAFs’ which they propose originate from a pool of perivascular cells^7^. The second most prominent population in this study were ‘matrix CAFs’, characterised by the expression of a wide variety of ECM and ECM-related genes that are proposed to originate from the activation of resident fibroblasts.

In this study we have focussed on the fibroblast receptor Endo180. Previously it has been reported that crossing an Endo180^−/−^ mouse with the MMTV-PyMT mouse resulted in reduction in primary mammary tumour burden^27^. However, the effects on metastatic colonisation were not investigated and little insight was provided as to the underlying mechanisms. Here we report that Endo180 expression is relatively low in the ‘vascular CAF’ subset^7^ and lymph node fibroblasts (bioRxiv doi.org/10.1101/467225), but instead represents one of the 30 core genes marking the population of ‘matrix CAFs’. Interestingly, a comparative analysis on mice with a genetic deletion in a vascular CAF receptor endosialin (*Cd248*) revealed a strikingly different phenotype compared to the Endo180 knockout mice. Whereas the reduced metastasis observed in the endosialin^−/−^ mice was due to a defect to tumour cell dissemination^43^, the Endo180^−/−^ mice display a defect in tumour growth and metastatic colonisation (Fig. 2 and 3) supporting the contention of different functional roles for different fibroblast subsets.

Although Endo180 is expressed at low level on stromal fibroblasts in normal adult tissues it is notable that non-tumour bearing Endo180 knockout mice have no notable phenotypic abnormalities and are born at the expected Mendelian frequency^16, 17^. However, a combined knockout of both Endo180 and the membrane type 1-matrix metalloproteinase (MT1-MMP), a major component of extracellular collagen degradation, resulted in accelerated postnatal lethality and severe impairment in bone growth^44^. These data, in agreement with our current and other studies^27^, suggest that loss of Endo180 in normal tissues can be compensated for, but that under conditions where matrix remodelling is compromised such as by loss of MT1-MMP or in the altered conditions of tumour microenvironment, a requirement for functional Endo180 is revealed. However, these previous studies have not addressed the mechanism by which loss of Endo180 expression impairs postnatal bone growth or tumour development and have focussed on the roles of Endo180 as an ECM remodelling receptor via its ability to internalise collagen for intracellular degradation. Although there is no question that Endo180 can function as a collagen internalisation receptor^15–19, 27, 45^, we demonstrate here that Endo180-deficient fibroblasts show impaired contractility when plated onto matrix-coated hydrogels but that this is not collagen dependent as an equivalent phenotype is observed when fibroblasts are plated onto fibronectin-coated hydrogels (Fig. 5; Supplementary Fig. 6). Similarly, we have previously reported that the defect in cell migration and contractile signalling associated with Endo180 depletion is not collagen dependent^30^. These data point to a CAF intrinsic defect in the Endo180^−/−^ tumour stroma and indeed we demonstrate here that Endo180 depletion is associated with increased fibroblast apoptosis when plated in 3D low serum culture and decreased fibroblast viability in 3D tumour-spheroid co-culture. Importantly, this is echoed by a reduced fibroblast content in the tumour stroma of Endo180^−/−^ mice and a reduction in collagen content in the tumours (Fig. 6). Interestingly, Wagenaar-Miller and colleagues reported that the Endo180 single knockout mice showed no difference in the number of apoptotic cells in the developing bones compared to wildtype mice. However, although an increased number of apoptotic cells were detected in the single MT1-MMP knockout mice there was significantly increased apoptosis and a corresponding decrease in cell proliferation in the double knockout compared to either single knockout^44^. Further, up-regulation of Endo180 expression has been shown for pathological conditions such as liver fibrosis induced by CCl_4_^46^, lung fibrosis induced by bleomycin^47^ and during wound healing^48, 49^. During wound healing, myofibroblasts are characterised by increased expression of αSMA^50^ but an inverse correlation between proliferation and ECM deposition^51^. At the end of the normal wound healing process^52^ or during the resolution of fibrosis^53^, myofibroblasts undergo cell death. However, evasion of apoptosis can lead to fibrosis via persistent myofibroblast activation and contraction-induced latent TGFβ activation^52, 54^. Furthermore, it has been demonstrated that reduction in mechanical stiffness in the matrix leads to an increase in fibroblast apoptosis^55, 56^. Hence, we suggest that a parallel situation may occur in tumours and that Endo180-deficient fibroblasts, which have reduced contractility and responsiveness to TGFβ (Fig. 5), will be cleared from the stromal compartment whilst in wildtype CAFs, Endo180 expression serves to maintain cell viability in pathological conditions.

Finally, we addressed how tumours might respond to a defective microenvironment by selecting for a tumour subline, D2A1-m12, by serial *in vivo* passage in Endo180^−/−^ mice. Compared to the parental D2A1 cells, D2A1-m12 cells were characterised by a striking upregulation of ECM and ECM-associated genes, as well as by increased αSMA (*Acta2*) expression. Together, these data suggest that D2A1-m12 subline is selected for a subclones(s) enriched for specific mesenchymal properties required to overcome the stromal deficit in the Endo180^−/−^ mice; providing further evidence for the critical role played by the tumour microenvironment in driving tumour evolution. Most importantly, when considering strategies for targeting the tumour stroma, these findings highlight the significance of limiting stromal activation early in the metastatic process.

## Methods

### Reagents and cells

Antibodies, and the dilutions used, are detailed in Supplementary Table 2. CN03 (Rho Activator-II, Cytoskeleton, Inc. #CN03-B). TGFβ1 (R&D Systems). 4T1, 4T07, MRC5, MDA-MB-453 and NIH-3T3 cells were from Isacke laboratory stocks. D2A1 cells were from Ann Chambers laboratory stocks^57^. The generation of the metastatic D2A1-m2 subline has been described previously^34^. The D2A1-m12 subline was generated as described for the D2A1-m2 subline except that the *in vivo* selection was in BALB/c Endo180^−/−^ mice. In brief, parental D2A1 cells were inoculated into the 4^th^ mammary fat pad of a BALB/c Endo180^−/−^ mouse, the lungs removed at necropsy, dissociated and placed into culture. Tumour cells that grew out were expanded and inoculated into the tail vein of a recipient Endo180^−/−^ mouse and 11-13 days later, lungs were removed at necropsy. In total three rounds of intravenous inoculation were performed. E0771 cells were purchased from CH3 BioSystems. Immortalised NF#1 mouse mammary gland fibroblasts were obtained from Fernando Calvo and have been described previously^58, 59^. 4T1, D2A1 and E0771 cells were luciferase transduced with lentiviral expression particles containing a firefly luciferase gene and a blasticidin-resistance gene (Amsbio, LVP326). Where indicated, 4T1-Luc cells were transduced with lentiviral particles expressing H2B-mRFP as previously described^58^ as well as D2A1, D2A1-m12 and MDA-MB-453 cells with a luciferase-mCherry vector. mCherry+/RFP+ cells were enriched by fluorescence-activated cell sorting (FACS). CAFs were isolated from a 4T1-tumour-bearing BALB/c Ub-GFP mouse (BALB/c mice expressing GFP under the human ubiquitin C promoter^60^. In brief, 1 × 10^4^ 4T1-Luc-RFP cells were injected into the 4^th^ mammary fat pad of a BALB/c Ub-GFP mouse. After 29 days the primary tumour was resected and homogenized using a McIlwain Tissue Chopper (Campden Instruments) and digested in L-15 medium containing 3 mg mL^−1^ collagenase type I at 37°C for 1 h, followed by digestion with 0.025 mg mL^−1^ DNase (Sigma) at 37 °C for 5 minutes. After erythrocyte lysis using Red Blood Cell Lysis Buffer (Sigma), single cells were re-suspended at a density of 1-2 × 10^7^ and subjected to magnetic sorting to exclude immune cells using rat-anti-mouse CD45 (R&D, MAB114) and CD24 (eBioscience, clone M1/69) magnetic sheep-anti-rat Ig Dynabeads (Invitrogen, #11035). FACS for FITC+/RFP-/DAPI-was used to obtain GFP+ve CAFs and to exclude RFP+ve tumour cells and dead cells, respectively. Cells were maintained in Dulbecco’s modified Eagle’s medium (DMEM) plus 10% foetal bovine serum (FBS), 1% penicillin/streptomycin and 1% ITS (insulin-transferrin-selenium; Invitrogen). All cells were routinely subject to mycoplasma testing.

shRNA Endo180 knock-down clones from 3T3 and CAFs were generated using lentiviral particles (Sigma, Supplementary Table 3). For siRNA knockdown, two non-targeting controls and two Endo180 targeting oligonucleotides (Supplementary Table 4) were used either separately or as pools in a 1:1 ratio.

### *In vitro* studies

#### Conditioned media (CM)

CM was generated by culturing fibroblasts or tumour cells at a 70-80% confluence. Cells were washed 2 times and then cultured in serum-free DMEM. After 24 hours, CM was collected, centrifuged at 300*g* and filtered through a 0.2 μm pore filter. Where indicated, CM was supplemented post-collection with FBS for long-term assays.

#### Cell proliferation/viability assays

1 × 10^3^ cells/well were seeded into 96-well plates. Cell viability was quantified either by CellTiter-Glo (Promega) at the indicated time points or by time-lapse imaging and quantification of cell confluence using the Live-Cell Analysis System IncuCyte (EssenBioscience).

#### Colony formation assay

50 tumour cells were seeded into a 6-well plate and either co-cultured with 5000 fibroblasts, treated with fibroblast conditioned media or plated onto fibroblast-derived ECM. Tumour colonies were stained 7 days later with crystal violet (Sigma). Plates were scanned using the GelCount (Oxford Optronix) and image analysis performed using GelCount software and FIJI.

#### Spheroid assays

Tumour cells and fibroblasts were mixed in a 1:9, 1:4 or 1:2 ratio and seeded into ultra-low adherence 96-well U-bottom plates (Corning). Spheroid growth was monitored using the Celigo Image Cytometer (Nexcelom Bioscience) and cell viability was monitored with CellTiter-Glo at the indicated time points. To ensure proper lysis of the spheroids, the incubation time with the CellTiter-Glo reagent was extended from 10 to 30 minutes before recording luminescence. As indicated cases, spheroids containing D2A1 cells and GFP-fibroblasts were dissociated using Accumax solution (Sigma) and cell numbers were quantified using the Celigo Image Cytometer (Nexcelom Bioscience) and a fluorescent Countess II FL Automated Cell Counter (Invitrogen).

#### Generation of extracellular matrices (ECM)

24-well plates were coated with 0.2% gelatin (Sigma, G1393) and cross-linked with 1% glutaraldehyde/PBS. Cross-linking was quenched with 1 M glycine/PBS. Fibroblasts were cultured in DMEM or tumour cell conditioned media supplemented with 2% FBS and 50 μg mL^−1^ ascorbic acid ascorbic acid (Sigma, A4403) on gelatin-coated plates. The medium was changed every 2 days for 7 days and fibroblasts were lysed using pre-warmed extraction buffer per well (20 mM NH_4_OH, 0.5% Triton X-100 in PBS). Residual DNA was digested with 10 μg mL^−1^ DNase I (Sigma). Matrices were either fixed with 4% paraformaldehyde and stained for fibronectin. Confocal z-stacks were acquired using the ImageXpress Micro Confocal High-Content Analysis System (Molecular Devices) equipped with a 60 μm pinhole spinning-disk, and a plan apo λ 10×/0.45 NA objective (Nikon). Z-stacks were maximally projected and analysed with ImageJ1.52p and the “TWOMBLI” plugin (bioRxiv doi.org/10.1101/867507), to quantify matrix density and (an-)isotropy of fibres. Data shown are matrix density, calculated as (100 − HDMvalue × 100) to convert from high density matrix background to % fibre density. End and branching points were normalised by dividing the raw value by the total length of the fibres. Alternatively, matrices in 24-well plates were extensively washed with PBS before tumour cells were seeded for a colony formation assay (50 cells per well) or IncuCyte imaging as described above (4000 cells per well).

#### Cell shape analysis

1 × 10^3^ cells/well were seeded into a ViewPlate-96 Black with optically clear bottom (PerkinElmer) or Softwell 24, Easy Coat plates (Cell guidance system) with the indicated stiffness. Plates were coated with 5 g cm^−2^ rat-tail collagen I (Corning, #354236) or fibronectin (R&D, 1918-FN-02M). After 24 hours, cells were fixed in 4% paraformaldehyde, permeabilised with 0.5% Triton X-100 prior to staining with DAPI and Alexa488- or Alexa555-labeled phalloidin. Automated image acquisition was performed on an Operetta high content imaging system (Perkin Elmer). Cell shapes were analysed using basic algorithms in the Harmony high content analysis software package (Perkin-Elmer). Cells were initially defined using the DAPI channel to identify the nucleus and the cytoplasm was segmented using the Alexa488/Alexa555 channel. After this, the Harmony software allows the extraction of cell shape parameters, including “cell roundness”.

#### Fibroblast contraction assays

7 × 10^4^ − 1 × 10^5^ fibroblasts were embedded in rat tail collagen I (Corning; final concentration 2 mg mL^−1^) in 24-well plates and incubated at 37°C. After the gels were set, media or conditioned media was added, as indicated. 24-72 hours later, plates were scanned and the contracted gel area was measured using FIJI.

#### Invasion assays

1 × 10^3^ fibroblasts were seeded into ultra-low adherence 96-well U bottom plates. 24 hours later, fibroblast ‘spheroids’ were embedded into 100 μL 2 mg mL^−1^ collagen (rat tail collagen, Corning). Invasion into collagen was imaged at the indicated time points and analysed using FIJI.

#### Apoptosis assay

5 × 10^4^ cells/well were plated into either a tissue culture or low adherence 6-well plates. 24 hours after seeding, cells were stained with the Annexin V-APC/ PI Apoptosis Detection Kit (eBioscience) and analysed using a BD Biosciences LSRII flow cytometer with FACSDIVA and FlowJo software.

#### Western blotting

Cells were lysed in RIPA lysis buffer supplemented with phosphatase inhibitors (PhosStop, Roche) and protease inhibitors (Complete, Roche). Cell lysates were subject to immunoblotting with the Bio-Rad Western blot system according to manufacturer’s protocol.

### *In vivo* procedures

All animal work was carried out under UK Home Office Project Licenses 70/7413 and P6AB1448A granted under the Animals (Scientific Procedures) Act 1986 (Establishment Licence, X702B0E74 70/2902) and was approved by the “Animal Welfare and Ethical Review Body” at The Institute of Cancer Research (ICR). Mice with a genetic deletion in Endo180 (*Mrc2*)^17^ were backcrossed for at least six generations with either BALB/c or C57BL/6 (Charles River) mice. Genotypes were confirmed by PCR. Endo180^−/−^ colonies were maintained at the ICR. Aged-matched 6-8 week-old female BALB/c or C57BL/6 mice were purchased from Charles River. All mice were housed in individually ventilated cages, monitored daily by ICR Biological Services Unit staff and had food and water *ad libitum*. In all cases, experiments were terminated if the primary tumour reached a maximum allowable diameter of 17 mm or if a mouse showed signs of ill health.

#### Spontaneous metastasis assays

1-5 × 10^4^ 4T1 or D2A1 cells were injected into the 4^th^ mammary fat pad under general anaesthesia. Tumour growth was measured twice a week using callipers up to a maximum diameter of 17 mm. Tumour volume^61^ was calculated as 0.5236 × [(width + length)/2]^3^. *Intravenous inoculation*: 1 × 10^5^ 4T1-Luc, 2.5 × 10^5^ 4T07, 4 × 10^5^ D2A1, D2A1-m2-Luc, or 4 × 10^5^ E0771-Luc cells were injected into the lateral tail vein of mice. Mice were sacrificed when the first animal showed signs of ill health. *Intrasplenic inoculation*: 1 × 10^5^ 4T1-Luc cells were inoculated into the spleen parenchyma of BALB/c mice under general anaesthesia.

Unless otherwise stated, primary tumours and lungs were weighed at necropsy. For IVIS imaging, mice were injected intraperitoneally with 150 mg kg^−1^ D-luciferin (Caliper Life Sciences) in 100 μL and mice imaged *in vivo* using an IVIS imaging chamber (IVIS Illumina II). Luminescence measurements (photons/second/cm^2^) were acquired over 1 minute and analysed using the Living Image software (PerkinElmer) using a constant size region of interest over the tissues. Alternatively, 5 minutes after D-luciferin injection, dissected lungs and/or livers were imaged *ex vivo*.

##### Quantification of metastatic burden and immunohistochemistry

3-4 μm lung and/or liver FFPE sections, approximately 150 μm apart, were cut and stained with haematoxylin and eosin (H&E). Sections were scanned using the NanoZoomer Digital Pathology and file names blinded. Total number of individual nodules was counted manually in 3 sections, per animal. Lung metastatic area was quantified as the mean % tumour area per lung section or by counting macroscopic tumour nodules counted manually. FFPE sections of primary mouse tumours were stained with αSMA and detection was achieved with the VectaStain ABS system. Stained sections were scanned on the NanoZoomer Digital Pathology (Hamamatsu). Images were exported and analysed with ImageJ and NDPITools ImageJ plugin^62^. HRP staining was analysed in ImageJ from ≥ 6 random fields of view per tumour section, avoiding areas of necrosis. HRP images were colour deconvoluted using the ImageJ H DAB vector and converted into 8-bit images and the % area quantified (threshold, 0-130). Collagen content in Masson’s trichrome stained sections was analysed similar to HRP staining, using the ImageJ Masson’s trichrome Vector.

Endo180 immunohistochemical staining in human samples was performed using anti-Endo180 mAb 39.10^25^.

### RT-qPCR

RNA was isolated using Qiagen RNeasy kit and cDNA was generated using the QuantiTect reverse transcription kit (Qiagen) according to the manufacturer’s instruction. qPCR was performed with human or mouse Taqman Gene Expression Assay probes on an ABI Prism 7900HT and relative quantification was performed using QuantStudio Real-time PCR software. Each reaction was performed in triplicate. Relative expression levels were normalised to *B2m/B2M* or *Gapdh/GAPDH* endogenous control, with a confidence interval of 95% for all assays. For co-culture Endo180 (*MRC2*) expression analysis, MRC5 lung fibroblasts were directly co-cultured with either D2A1-RFP or MDA-MB-453-RFP cells for 48 hours. Cells were sorted by FACS for RFP-ve fibroblasts, which were further subjected to RNA isolation and RT-qPCR analysis.

### RNA-Seq analysis

RNA from D2A1 and D2A1-m12 cells (n=3 per line) was extracted using the RNeasy kit according to the manufacturer’s instructions. Quality and quantity of RNA were assessed using a Qubit and Bioanalyzer. NEBNEXT Ultra II Directional RNA and polyA RNA selection kits were used to generate libraries, which were combined and sequenced at PR 100 cycles on a NovaSeq flowcell. All samples were run to achieve ~50 million clusters. RNA-Seq generated 11.4 to 53.1 million reads per sample. FastQC (v0.11.4) was used to evaluate the library quality. Paired-end reads (100 base pair long) were aligned to the mouse reference genome GRCm38, using STAR v2.5.1b^63^ with --quantMode GeneCounts and --twopassMode Basic alignment settings. Annotation file used for feature quantifications was downloaded from GENCODE (v17) in GTF file format. Post alignment quality control was performed using RseQC (v2.6.3)^64^.

Differential mRNA abundance analysis was performed using edgeR package (v3.22.5)^65^ in R (v3.5.0) using the model *~0 + group.* Genes with low expression were filtered out by retaining genes with count-per-million (CPM) counts > 1 in at least two samples. Samples were normalised using edgeR’s TMM (trimmed mean of M-values) method and differential expression was determined using quasi-likelihood (QL) F-test. Results were further annotated using ENSEMBL gene annotations from R package org.Mm.eg.db (v3.6.0). Genes with |log_2_FC| > 1 and adjusted *P* value ≤ 0.05 were considered statistically significant.

Fast Gene Set Enrichment Analysis (fGSEA) was performed with R package fgsea (v1.8.0) using mouse gene sets (Biocarta, NetPath, Panther) downloaded from GSKB (Gene Set Knowledgebase in Mouse) database, and the mouse gene sets obtained from the Matrisome Project (matrisomeproject.mit.edu). Genes were ranked as: – log_10_(P) × sgn(log_2_FC). We applied a minimum gene set size of 10 genes and performed the analysis using 10,000 permutations. We considered pathways with normalised enrichment score (NES) > 0 and *P* value < 0.05 to be upregulated and pathways with NES < 0 and *P* value < 0.05 to be downregulated.

### Analysis of human samples and datasets

FFPE sections were stained with 40 μg/mL anti-Endo180 mAb 39.10^25^ for 30 min at room temperature and detection was achieved with the Dako Envision/HRP system.

### Bioinformatics analysis

Series matrix files for TCGA 522 primary breast cancer samples were downloaded from [https://tcga-data.nci.nih.gov/docs/publications/brca_2012/]. GSE39397^31^, GSE37614^66^, GSE41678^32^, GSE111229^7^ were downloaded from the Gene Expression Omnibus (GEO) site. NKI295^67^ from http://ccb.nki.nl/data/. Graphs show gene expression or median centred gene expression levels. Cell-types in GSE39397 were defined as EpCAM+ve, epithelial cells, CD45+ve leukocytes, CD31+ve endothelial cells and FAP+ve fibroblasts according to original publication^31^. Intrinsic molecular subtypes (luminal A, luminal B, HER2-enriched, basal-like) and normal versus adjacent data were retrieved from the supplemental tables of the corresponding publications. In boxplots, box indicates the ends of the 1st and 3rd quartiles, bar indicates median, whiskers indicated 1.5 IQR (interquartile range), and dots indicate outliers of gene expression.

For association of Endo180 (*MRC2)* gene expression in TCGA and subsets (luminal A, luminal B, HER2-enriched, basal-like) and NKI dataset with fibroblast TGFβ response signature (F-TBRS), the gene set was retrieved from http://dx.doi.org/10.1016/j.ccr.2012.08.013^31^. There were 286 probes representing 171 unique annotated genes. In the TCGA dataset 151 out of the 171 F-TBRS genes were matched. Average gene expressions of the 151 genes from TCGA dataset was used as signature score for each tumour.

Endo180 (Mrc2) gene expression level log_2_(RPKM+1) from single cell sequencing of CAFs isolated from MMTV-PyMT tumours^7^ are clustered according to identified populations in the original publication (vascular CAFs, matrix CAFs, developmental CAFs and actively cycling CAFs).

### Statistics

Statistics were performed using GraphPad Prism 8. Unless otherwise indicated, all comparisons between two groups were made using two-tailed, unpaired Student’s *t*-test. If the analysis did not pass normality test (Shapiro-Wilk test) groups were analysed by Mann Whitney *U* test. If more than 2 groups were compared One-way ANOVA analysis was performed with Tukey test for multiple comparison. Where multiple groups with a second variable, e.g. over time, were compared, a two-way ANOVA followed by Sidak post-hoc testing was performed. *, *P* < 0.05; ** *P* < 0.01; ***, *P* < 0.001.

### Data availability

Raw RNA-Seq and whole exome sequencing (WES) data will be made available via ENA.

## Acknowledgements

This work was funded by Breast Cancer Now (CTR-Q4-Y3), working in partnership with Walk the Walk (CMI), Worldwide Cancer Research (15-0355, CMI and UJ) and a Schrödinger fellowship of the Austrian Science Fund (FWF) (J3434-B13, UJ). We acknowledge NHS funding to the NIHR Biomedical Research Centre at the Royal Marsden and the ICR. We would like to thank Adam Mills for the whole exome sequencing analysis, Kristian Pietras and Jonas Sjölund for the data analysis and images in Figure 1h, Miriam Melake for help with the matrix imaging, Gernot Walko for his imaging expertise and useful discussions, and the following facilities at the ICR; Tumour Profiling Unit, Biological Services Unit, FACS and Light Microscopy Facility and the Breast Cancer Now Histopathology Facility. The results published here are in whole or part based upon data generated by the TCGA Research Network (http://cancergenome.nih.gov/).

## Author Contributions

Conception and design of the work (UJ, CMI)

Acquisition, analysis, interpretation of data (UJ, AvW, LJ, JA, DV, QG, SH, MI, CMI)

Preparation of the manuscript (UJ, SH, MI, CMI with input from all other authors)

## Competing Interests statement

The authors have no conflicts of interest to declare

## Notes

### Competing Interest Statement

The authors have declared no competing interest.

## References

1. Harper, J. & Sainson, R.C. Regulation of the anti-tumour immune response by cancer-associated fibroblasts. Semin Cancer Biol 25, 69–77 (2014).

2. Kalluri, R. The biology and function of fibroblasts in cancer. Nature reviews. Cancer 16, 582–598 (2016).

3. Valkenburg, K.C., de Groot, A.E. & Pienta, K.J. Targeting the tumour stroma to improve cancer therapy. Nat Rev Clin Oncol 15, 366–381 (2018).

4. Zambirinis, C.P. & Miller, G. Cancer Manipulation of Host Physiology: Lessons from Pancreatic Cancer. Trends Mol Med 23, 465–481 (2017).

5. Sahai, E. et al. A framework for advancing our understanding of cancer-associated fibroblasts. Nature reviews. Cancer (2020).

6. LeBleu, V.S. & Kalluri, R. A peek into cancer-associated fibroblasts: origins, functions and translational impact. Dis Model Mech 11(2018).

7. Bartoschek, M. et al. Spatially and functionally distinct subclasses of breast cancer-associated fibroblasts revealed by single cell RNA sequencing. Nat Commun 9, 5150 (2018).

8. Brechbuhl, H.M. et al. Fibroblast Subtypes Regulate Responsiveness of Luminal Breast Cancer to Estrogen. Clinical cancer research: an official journal of the American Association for Cancer Research 23, 1710–1721 (2017).

9. Costa, A. et al. Fibroblast Heterogeneity and Immunosuppressive Environment in Human Breast Cancer. Cancer cell 33, 463–479 e410 (2018).

10. Neuzillet, C. et al. Inter- and intra-tumoural heterogeneity in cancer-associated fibroblasts of human pancreatic ductal adenocarcinoma. J Pathol 248, 51–65 (2019).

11. Ohlund, D. et al. Distinct populations of inflammatory fibroblasts and myofibroblasts in pancreatic cancer. The Journal of experimental medicine 214, 579–596 (2017).

12. Puram, S.V. et al. Single-Cell Transcriptomic Analysis of Primary and Metastatic Tumor Ecosystems in Head and Neck Cancer. Cell 171, 1611–1624 e1624 (2017).

13. Su, S. et al. CD10(+)GPR77(+) Cancer-Associated Fibroblasts Promote Cancer Formation and Chemoresistance by Sustaining Cancer Stemness. Cell 172, 841–856 e816 (2018).

14. Zhao, Q. et al. Single-Cell Transcriptome Analyses Reveal Endothelial Cell Heterogeneity in Tumors and Changes following Antiangiogenic Treatment. Cancer Res 78, 2370–2382 (2018).

15. Wienke, D., MacFadyen, J.R. & Isacke, C.M. Identification and characterization of the endocytic transmembrane glycoprotein Endo180 as a novel collagen receptor. Mol Biol Cell 14, 3592–3604 (2003).

16. Engelholm, L.H. et al. uPARAP/Endo180 is essential for cellular uptake of collagen and promotes fibroblast collagen adhesion. The Journal of cell biology 160, 1009–1015 (2003).

17. East, L. et al. A targeted deletion in the endocytic receptor gene Endo180 results in a defect in collagen uptake. EMBO reports 4, 710–716 (2003).

18. Kjøller, L. et al. uPARAP/endo180 directs lysosomal delivery and degradation of collagen IV. Experimental cell research 293, 106–116 (2004).

19. Jurgensen, H.J. et al. Complex determinants in specific members of the mannose receptor family govern collagen endocytosis. The Journal of biological chemistry 289, 7935–7947 (2014).

20. East, L., Rushton, S., Taylor, M.E. & Isacke, C.M. Characterization of sugar binding by the mannose receptor family member, Endo180. Journal of Biological Chemistry 277, 50469–50475 (2002).

21. Howard, M.J. & Isacke, C.M. The C-type lectin receptor Endo180 displays internalization and recycling properties distinct from other members of the mannose receptor family. Journal of Biological Chemistry 277, 32320–32331 (2002).

22. Isacke, C., Van der Geer, P., Hunter, T. & Trowbridge, I. p180, a novel recycling transmembrane glycoprotein with restricted cell type expression. Molecular and cellular biology 10, 2606–2618 (1990).

23. Engelholm, L.H. et al. Targeting a novel bone degradation pathway in primary bone cancer by inactivation of the collagen receptor uPARAP/Endo180. J Pathol 238, 120–133 (2016).

24. Huijbers, I.J. et al. A role for fibrillar collagen deposition and the collagen internalization receptor endo180 in glioma invasion. PloS one 5, e9808 (2010).

25. Wienke, D. et al. The collagen receptor Endo180 (CD280) Is expressed on basal-like breast tumor cells and promotes tumor growth in vivo. Cancer Research 67, 10230–10240 (2007).

26. Schnack Nielsen, B. et al. Urokinase receptor-associated protein (uPARAP) is expressed in connection with malignant as well as benign lesions of the human breast and occurs in specific populations of stromal cells. Int J Cancer 98, 656–664 (2002).

27. Curino, A.C. et al. Intracellular collagen degradation mediated by uPARAP/Endo180 is a major pathway of extracellular matrix turnover during malignancy. J Cell Biol 169, 977–985 (2005).

28. Koikawa, K. et al. Pancreatic stellate cells reorganize matrix components and lead pancreatic cancer invasion via the function of Endo180. Cancer Lett 412, 143–154 (2018).

29. Sulek, J. et al. Increased expression of the collagen internalization receptor uPARAP/Endo180 in the stroma of head and neck cancer. J Histochem Cytochem 55, 347–353 (2007).

30. Sturge, J., Wienke, D. & Isacke, C.M. Endosomes generate localized Rho– ROCK–MLC2–based contractile signals via Endo180 to promote adhesion disassembly. The Journal of cell biology 175, 337–347 (2006).

31. Calon, A. et al. Dependency of colorectal cancer on a TGF-beta-driven program in stromal cells for metastasis initiation. Cancer cell 22, 571–584 (2012).

32. Rajaram, M., Li, J., Egeblad, M. & Powers, R.S. System-wide analysis reveals a complex network of tumor-fibroblast interactions involved in tumorigenicity. PLoS Genet 9, e1003789 (2013).

33. Calon, A. et al. Stromal gene expression defines poor-prognosis subtypes in colorectal cancer. Nat Genet 47, 320–329 (2015).

34. Jungwirth, U. et al. Generation and characterisation of two D2A1 mammary cancer sublines to model spontaneous and experimental metastasis in a syngeneic BALB/c host. Dis Model Mech 11(2018).

35. Flatau, G. et al. Toxin-induced activation of the G protein p21 Rho by deamidation of glutamine. Nature 387, 729–733 (1997).

36. Schmidt, G. et al. Gln 63 of Rho is deamidated by Escherichia coli cytotoxic necrotizing factor-1. Nature 387, 725–729 (1997).

37. Merlo, L.M., Pepper, J.W., Reid, B.J. & Maley, C.C. Cancer as an evolutionary and ecological process. Nature reviews. Cancer 6, 924–935 (2006).

38. Naba, A. et al. The matrisome: in silico definition and in vivo characterization by proteomics of normal and tumor extracellular matrices. Mol Cell Proteomics 11, M111 014647 (2012).

39. Chen, X. & Song, E. Turning foes to friends: targeting cancer-associated fibroblasts. Nat Rev Drug Discov 18, 99–115 (2019).

40. Shiga, K. et al. Cancer-Associated Fibroblasts: Their Characteristics and Their Roles in Tumor Growth. Cancers (Basel) 7, 2443–2458 (2015).

41. Crisan, M. et al. A perivascular origin for mesenchymal stem cells in multiple human organs. Cell Stem Cell 3, 301–313 (2008).

42. Shi, Y., Du, L., Lin, L. & Wang, Y. Tumour-associated mesenchymal stem/stromal cells: emerging therapeutic targets. Nat Rev Drug Discov 16, 35–52 (2017).

43. Viski, C. et al. Endosialin-Expressing Pericytes Promote Metastatic Dissemination. Cancer Res 76, 5313–5325 (2016).

44. Wagenaar-Miller, R.A. et al. Complementary roles of intracellular and pericellular collagen degradation pathways in vivo. Mol Cell Biol 27, 6309–6322 (2007).

45. Madsen, D.H. et al. The non-phagocytic route of collagen uptake: a distinct degradation pathway. The Journal of biological chemistry 286, 26996–27010 (2011).

46. Madsen, D.H. et al. Endocytic collagen degradation: a novel mechanism involved in protection against liver fibrosis. The Journal of pathology (2012).

47. Jurgensen, H.J. et al. Immune regulation by fibroblasts in tissue injury depends on uPARAP-mediated uptake of collectins. J Cell Biol 218, 333–349 (2019).

48. Rohani, M.G. et al. uPARAP function in cutaneous wound repair. PloS one 9, e92660 (2014).

49. Honardoust, H.A. et al. Expression of Endo180 is spatially and temporally regulated during wound healing. Histopathology 49, 634–648 (2006).

50. Ronnov-Jessen, L. & Petersen, O.W. Induction of alpha-smooth muscle actin by transforming growth factor-beta 1 in quiescent human breast gland fibroblasts. Implications for myofibroblast generation in breast neoplasia. Lab Invest 68, 696–707 (1993).

51. Rognoni, E. et al. Fibroblast state switching orchestrates dermal maturation and wound healing. Mol Syst Biol 14, e8174 (2018).

52. Hinz, B. & Lagares, D. Evasion of apoptosis by myofibroblasts: a hallmark of fibrotic diseases. Nat Rev Rheumatol 16, 11–31 (2020).

53. Jun, J.I. & Lau, L.F. Resolution of organ fibrosis. J Clin Invest 128, 97–107 (2018).

54. Biernacka, A., Dobaczewski, M. & Frangogiannis, N.G. TGF-beta signaling in fibrosis. Growth Factors 29, 196–202 (2011).

55. Graf, R., Freyberg, M., Kaiser, D. & Friedl, P. Mechanosensitive induction of apoptosis in fibroblasts is regulated by thrombospondin-1 and integrin associated protein (CD47). Apoptosis 7, 493–498 (2002).

56. Tian, B., Lessan, K., Kahm, J., Kleidon, J. & Henke, C. beta 1 integrin regulates fibroblast viability during collagen matrix contraction through a phosphatidylinositol 3-kinase/Akt/protein kinase B signaling pathway. The Journal of biological chemistry 277, 24667–24675 (2002).

57. Morris, V.L., Tuck, A.B., Wilson, S.M., Percy, D. & Chambers, A.F. Tumor progression and metastasis in murine D2 hyperplastic alveolar nodule mammary tumor cell lines. Clinical & experimental metastasis 11, 103–112 (1993).

58. Avgustinova, A. et al. Tumour cell-derived Wnt7a recruits and activates fibroblasts to promote tumour aggressiveness. Nat Commun 7, 10305 (2016).

59. Calvo, F. et al. Mechanotransduction and YAP-dependent matrix remodelling is required for the generation and maintenance of cancer-associated fibroblasts. Nature cell biology 15, 637–646 (2013).

60. Schaefer, B.C., Schaefer, M.L., Kappler, J.W., Marrack, P. & Kedl, R.M. Observation of antigen-dependent CD8+ T-cell/ dendritic cell interactions in vivo. Cell Immunol 214, 110–122 (2001).

61. Janik, P., Briand, P. & Hartmann, N.R. The effect of estrone-progesterone treatment on cell proliferation kinetics of hormone-dependent GR mouse mammary tumors. Cancer Res 35, 3698–3704 (1975).

62. Deroulers, C. et al. Analyzing huge pathology images with open source software. Diagn Pathol 8, 92 (2013).

63. Dobin, A. et al. STAR: ultrafast universal RNA-seq aligner. Bioinformatics 29, 15–21 (2013).

64. Wang, L., Wang, S. & Li, W. RSeQC: quality control of RNA-seq experiments. Bioinformatics 28, 2184–2185 (2012).

65. Robinson, M.D., McCarthy, D.J. & Smyth, G.K. edgeR: a Bioconductor package for differential expression analysis of digital gene expression data. Bioinformatics 26, 139–140 (2010).

66. Tchou, J. et al. Human breast cancer associated fibroblasts exhibit subtype specific gene expression profiles. BMC Med Genomics 5, 39 (2012).

67. van de Vijver, M.J. et al. A gene-expression signature as a predictor of survival in breast cancer. N Engl J Med 347, 1999–2009 (2002).

