## Supplementary Material for "Impairment of a distinct cancer-associated fibroblast population limits tumour growth and metastasis"

**Supplementary methods**

**Supplementary Tables 1 - 4**

Supplementary Table 1a. | Top 30 upregulated pathways in D2A1-m12 cells compared to D2A1 cells

Supplementary Table 1b. | Top 30 downregulated pathways in D2A1-m12 cells compared to D2A1 cells

Supplementary Table 2 | Antibodies

Supplementary Table 3 | Mission shRNA lentiviral particles targeting Endo180 (*Mrc2*) (Sigma)

Supplementary Table 4 | ON-TARGET *plus* siRNA targeting Endo180 (*Mrc2*) (Dharmacon)

**Supplementary Figures 1 - 7**

Supplementary Fig. 1 | Fibroblast expression of Endo180 in solid tumours

Supplementary Fig. 2 | Endo180 promotes metastatic tumour growth in lungs and liver

Supplementary Fig. 3 | Expression of Endo180 in the brain

Supplementary Fig. 4 | Endo180 expression following siRNA- or shRNA-mediated knockdown

Supplementary Fig. 5 | Fibroblast-tumour cell co-culture assays

Supplementary Fig. 6 | Cell spreading on collagen or fibronectin-coated hydrogels

Supplementary Fig. 7 | Comparison of the parental D2A1 line and the D2A1-m12 subline

**Supplementary methods**

Whole exome sequencing

DNA was extracted from tumour cell lines or BALB/c mouse spleen tissue using the DNeasy Blood and Tissue kit. DNA samples were physically sheared to the desired size using a Covaris E Series instrument (Covaris). Paired-end multiplexed library preparation was performed by the Tumour Profiling Unit at the ICR using the SureSelect$^{XT}$ Library Prep and Capture System (Agilent Technologies) following the standard protocol workflow, before multiplex sequencing on a NovaSeq 6000 flow cell (Illumina).

Bioinformatics and statistical analyses were performed using custom R scripts and a bespoke DNA sequencing pipeline based on the Nextflow framework[1]. Whole exome sequencing FASTQ files were aligned to the mouse genome assembly build GRCm38 with the Burrows-Wheeler Aligner (BWA) v0.7.12[2]. BWA was run with default parameters utilising 5 threads. The resulting SAM file was converted to BAM and sorted with SAMtools v1.5[3]. Duplicate reads were removed from the sample files with the Picard v2.8.1 suite of tools (http://broadinstitute.github.io/picard/). Insert size and coverage metrics were also calculated with Picard. Lastly, base scores were recalibrated with the Genome Analysis Toolkit (GATK) v4.0.3.0[4] according to the GATK Best Practices pipeline.

Copy number analysis (CNA) was carried out with CNVkit v0.9.3[5], utilising the batch command. A cancer-free sample taken from BALB/c mouse spleen tissue was used as a normal reference for each cell line sample. Copy number $\log_2$ ratios from CNVkit were used to create CNA plots in R statistical programming language v3.5.0 (R Core Team, 2018).

LoFreq[6] was used to call somatic variants. In CNA, the same healthy BALB/c sample was used as a normal reference. Common variants were removed before annotating the remaining variants with ANNOVAR (01/02/2016 release)[7]. The output from ANNOVAR was further processed in R for comparisons between samples and visualisations.

### Supplementary Tables

**Supplementary Table 1a. | Top 30 upregulated pathways in D2A1-m12 cells compared to D2A1 cells**

| Pathway | NES | FDR |
|---|---|---|
| mouse.corematrisome.collagens | 1.886 | 0.038 |
| mouse.matrisome.associated.ecm.affiliated | 1.869 | 0.019 |
| mouse.corematrisome.glycoproteins | 1.836 | 0.024 |
| BIOCARTA_MM_CASPASE_CASCADE_IN_APOPTOSIS | 1.793 | 0.058 |
| BIOCARTA_MM_THROMBIN_SIGNALING_AND_PROTEASE-ACTIVATED_RECEPTORS | 1.792 | 0.038 |
| BIOCARTA_MM_D4-GDI_SIGNALING_PATHWAY | 1.732 | 0.038 |
| PANTHER_MM_INTEGRIN_SIGNALLING_PATHWAY | 1.729 | 0.019 |
| BIOCARTA_MM_RHO_CELL_MOTILITY_SIGNALING_PATHWAY | 1.633 | 0.224 |
| mouse.matrisome.associated.regulators | 1.582 | 0.079 |
| PANTHER_MM_HETEROTRIMERIC_G-PROTEIN_SIGNALING_PATHWAY-GI_ALPHA_AND_GS_ALPHA_MEDIATED | 1.562 | 0.179 |
| PANTHER_MM_FAS_SIGNALING_PATHWAY | 1.548 | 0.345 |
| BIOCARTA_MM_PHOSPHOLIPIDS_AS_SIGNALLING_INTERMEDIARIES | 1.540 | 0.353 |
| BIOCARTA_MM_RAC_1_CELL_MOTILITY_SIGNALING_PATHWAY | 1.533 | 0.371 |
| BIOCARTA_MM_INTEGRIN_SIGNALING_PATHWAY | 1.525 | 0.345 |
| BIOCARTA_MM_RAS-INDEPENDENT_PATHWAY_IN_NK_CELL-MEDIATED_CYTOTOXICITY | 1.522 | 0.371 |
| PANTHER_MM_IONOTROPIC_GLUTAMATE_RECEPTOR_PATHWAY | 1.511 | 0.371 |
| BIOCARTA_MM_ROLES_OF_ARRESTIN-DEPENDENT_RECRUITMENT_OF_SRC_KINASES_IN_GPCR_SIGNALING | 1.493 | 0.393 |
| PANTHER_MM_INFLAMMATION_MEDIATED_BY_CHEMOKINE_AND_CYTOKINE_SIGNALING | 1.489 | 0.156 |
| BIOCARTA_MM_ROLE_OF_ARRESTINS_IN_THE_ACTIVATION_AND_TARGETING_OF_MAP_KINASES | 1.477 | 0.399 |
| BIOCARTA_MM_SPLICEOSOMAL_ASSEMBLY | 1.470 | 0.399 |
| BIOCARTA_MM_CCR3_SIGNALING_IN_EOSINOPHILS | 1.467 | 0.403 |
| BIOCARTA_MM_TGF_BETA_SIGNALING_PATHWAY | 1.464 | 0.399 |
| NETPATH_MM_ALPHA6_BETA4_INTEGRIN | 1.423 | 0.371 |
| NETPATH_MM_EGFR1_SIGNALING_PATHWAY | 1.415 | 0.038 |
| mouse.matrisome.associated.secretedfactors | 1.409 | 0.345 |
| PANTHER_MM_CYTOSKELETAL_REGULATION_BY_RHO_GTPASE | 1.408 | 0.393 |
| NETPATH_MM_B_CELL_RECEPTOR_SIGNALING_PATHWAY | 1.403 | 0.314 |
| PANTHER_MM_HETEROTRIMERIC_G-PROTEIN_SIGNALING_PATHWAY-GQ_ALPHA_AND_GO_ALPHA_MEDIATED | 1.382 | 0.421 |
| PANTHER_MM_HUNTINGTON_DISEASE | 1.377 | 0.371 |
| BIOCARTA_MM_MTOR_SIGNALING_PATHWAY | 1.357 | 0.602 |
| BIOCARTA_MM_TREFOIL_FACTORS_INITIATE_MUCOSAL_HEALING | 1.352 | 0.602 |

Pathways highlighted in yellow have FDR < 0.1. FDR, false discovery rate. NES, normalised enrichment score

| Supplementary Table 1b. \| Top 30 downregulated pathways in D2A1-m12 cells compared to D2A1 cells | | |
|---|---|---|
| Pathway | NES | FDR |
| BIOCARTA_MM_SYNAPTIC_PROTEINS_AT_THE_SYNAPTIC_JUNCTION | -1.606 | 0.345 |
| PANTHER_MM_SYNAPTIC_VESICLE_TRAFFICKING | -1.531 | 0.391 |
| PANTHER_MM_INSULIN_IGF_PATHWAY-PROTEIN_KINASE_B_SIGNALING_CASCADE | -1.527 | 0.371 |
| PANTHER_MM_METABOTROPIC_GLUTAMATE_RECEPTOR_GROUP_I_PATHWAY | -1.429 | 0.558 |
| BIOCARTA_MM_BONE_REMODELLING | -1.404 | 0.602 |
| BIOCARTA_MM_REGULATION_OF_EIF2 | -1.400 | 0.600 |
| PANTHER_MM_EGF_RECEPTOR_SIGNALING_PATHWAY | -1.321 | 0.393 |
| BIOCARTA_MM_TELOMERES_TELOMERASE_CELLULAR_AGING_AND_IMMORTALITY | -1.299 | 0.646 |
| PANTHER_MM_MUSCARINIC_ACETYLCHOLINE_RECEPTOR_1_AND_3_SIGNALING_PATHWAY | -1.272 | 0.614 |
| BIOCARTA_MM_TOLL-LIKE_RECEPTOR_PATHWAY | -1.271 | 0.602 |
| PANTHER_MM_P53_PATHWAY_FEEDBACK_LOOPS_2 | -1.269 | 0.602 |
| PANTHER_MM_CORTOCOTROPIN_RELEASING_FACTOR_RECEPTOR_SIGNALING_PATHWAY | -1.264 | 0.667 |
| PANTHER_MM_PI3_KINASE_PATHWAY | -1.259 | 0.602 |
| BIOCARTA_MM_CD40L_SIGNALING_PATHWAY | -1.248 | 0.704 |
| PANTHER_MM_PDGF_SIGNALING_PATHWAY | -1.222 | 0.568 |
| PANTHER_MM_HISTAMINE_H1_RECEPTOR_MEDIATED_SIGNALING_PATHWAY | -1.205 | 0.692 |
| PANTHER_MM_INTERLEUKIN_SIGNALING_PATHWAY | -1.188 | 0.656 |
| BIOCARTA_MM_INHIBITION_OF_CELLULAR_PROLIFERATION_BY_GLEEVEC | -1.186 | 0.757 |
| PANTHER_MM_INSULIN_IGF_PATHWAY-MITOGEN_ACTIVATED_PROTEIN_KINASE_KINASE_MAP_KINASE_CASCADE | -1.177 | 0.753 |
| BIOCARTA_MM_P53_SIGNALING_PATHWAY | -1.094 | 0.808 |
| PANTHER_MM_OXIDATIVE_STRESS_RESPONSE | -1.092 | 0.795 |
| BIOCARTA_MM_P38_MAPK_SIGNALING_PATHWAY_ | -1.087 | 0.795 |
| BIOCARTA_MM_KERATINOCYTE_DIFFERENTIATION | -1.087 | 0.795 |
| BIOCARTA_MM_SKELETAL_MUSCLE_HYPERTROPHY_IS_REGULATED_VIA_AKT_MTOR_PATHWAY | -1.083 | 0.808 |
| PANTHER_MM_HYPOXIA_RESPONSE_VIA_HIF_ACTIVATION | -1.067 | 0.808 |
| BIOCARTA_MM_THE_4-1BB-DEPENDENT_IMMUNE_RESPONSE | -1.052 | 0.827 |
| PANTHER_MM_OXYTOCIN_RECEPTOR_MEDIATED_SIGNALING_PATHWAY | -1.050 | 0.824 |
| PANTHER_MM_5HT2_TYPE_RECEPTOR_MEDIATED_SIGNALING_PATHWAY | -1.031 | 0.827 |
| PANTHER_MM_HISTAMINE_H2_RECEPTOR_MEDIATED_SIGNALING_PATHWAY | -1.023 | 0.872 |
| BIOCARTA_MM_NEUROPEPTIDES_VIP_AND_PACAP_INHIBIT_THE_APOPTOSIS_OF_ACTIVATED_T_CELLS | -0.987 | 0.922 |
| BIOCARTA_MM_CERAMIDE_SIGNALING_PATHWAY | -0.970 | 0.922 |
| FDR, false discovery rate. NES, normalised enrichment score | | |

**Supplementary Table 2 | Antibodies**

| Antibody | Species | Source (catalogue number) | Assays | Dilution |
|---|---|---|---|---|
| αSMA | mouse | Sigma (clone 1A4) | IHC | 1:1,000 |
| CD24 | rat | eBioscience (M1/69) | FACS | 4 µL/1x$10^7$cells/mL |
| CD45 | rat | R&D Systems (MAB114) | FACS | 4 µL/1x$10^7$cells/mL |
| Endo180 (human) | mouse | mAb 39.10 (in house)[8] | IHC | 1:1,000 |
| Endo180 (mouse) | sheep | R&D Systems (AF4789) | WB | 1 µg/mL |
| Fibronectin | rabbit | Dako (A0245) | IF | 1:2,000 |
| MLC | rabbit | Cell Signaling (3672) | WB | 1:1,000 |
| p(ser19)-MLC | mouse | Cell Signaling (3675) | WB | 1:1,000 |
| Alexa488-phalloidin | N/A | Molecular Probes (A12379) | IF | 1:500 |
| Alexa555-phalloidin | N/A | Molecular Probes (A34055) | IF | 1:500 |
| DAPI | N/A | Molecular Probes (D1306) | IF | 1:10,000 |
| IgG-HRP-anti-mouse | donkey | Santa Cruz (sc-2314) | WB | 1:10,000 |
| IgG-HRP-anti-rabbit | goat | Santa Cruz (sc-2004) | WB | 1:10,000 |
| IgG-HRP-anti-sheep | rabbit | Santa Cruz (sc-2770) | WB | 1:10,000 |

FACS, fluorescence activated cell sorting; IF, immunofluorescence; IHC, immunohistochemistry; WB, western blot.

| **Supplementary Table 3 \| Mission shRNA lentiviral particles targeting Endo180 (*Mrc2*) (Sigma)** | | |
|---|---|---|
| **Clone ID** | **Gene target** | **NM ID** |
| SHC002V | NTC | N/A |
| TRCN00001239-25 | *Mrc2* | NM_008626.3 |
| TRCN00001239-27 | *Mrc2* | NM_008626.3 |
| TRCN00001239-28 | *Mrc2* | NM_008626.3 |

| **Supplementary Table 4 \| ON-TARGET *plus* siRNA targeting Endo180 (*Mrc2*) (Dharmacon)** | | |
|---|---|---|
| **ID** | **Gene target** | **NM ID** |
| D-001810-01 | NTC | N/A |
| D-001810-04 | NTC | N/A |
| J-040940-09 | *Mrc2* | NM_008626.3 |
| J-040940-12 | *Mrc2* | NM_008626.3 |

### Supplementary Figure 1

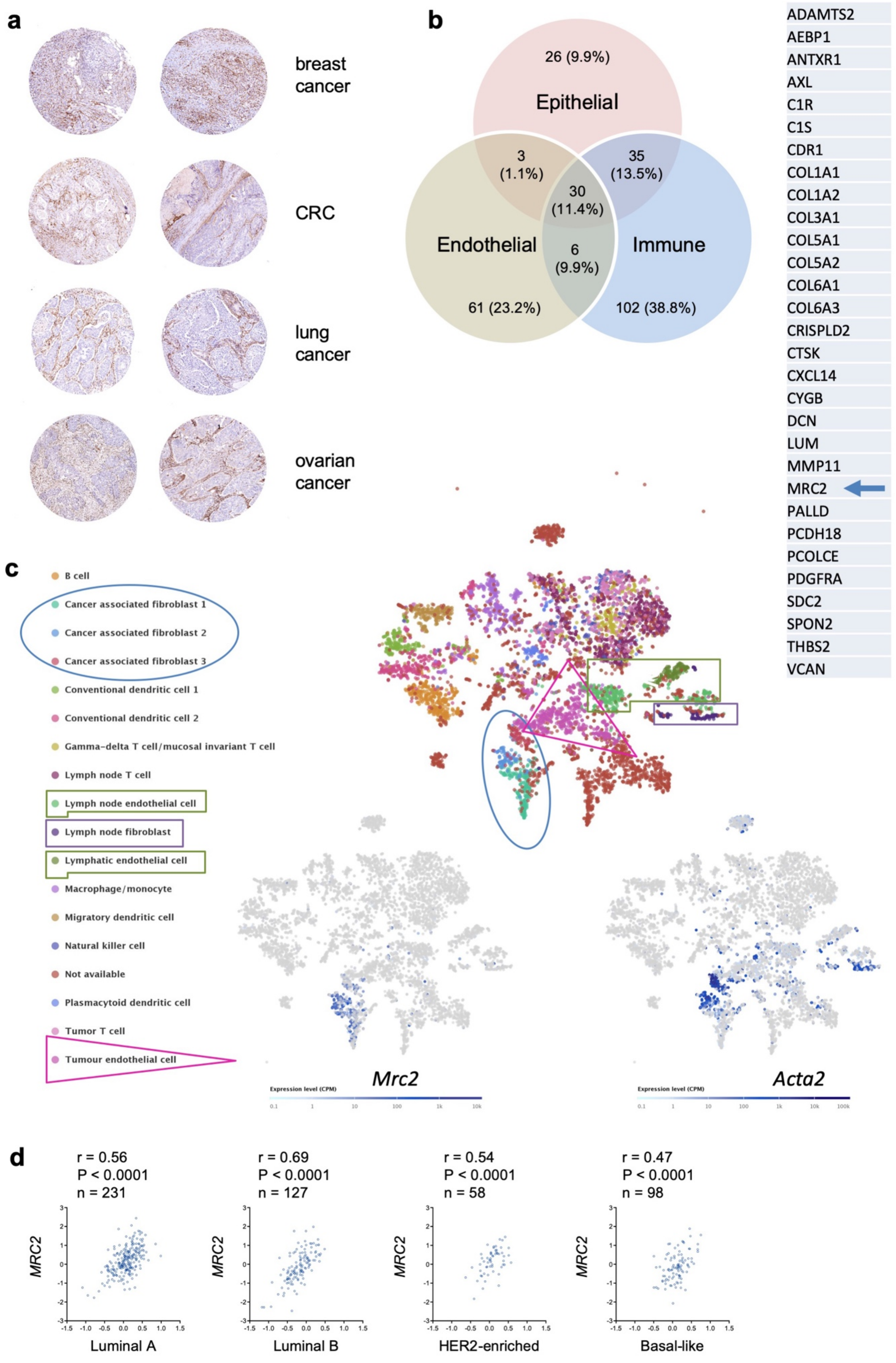

**Supplementary Fig. 1 | Fibroblast expression of Endo180 in solid tumours**
**a** Images taken from The Human Protein Atlas (www.proteinatlas.org) of breast, colorectal lung and ovarian tumours stained for Endo180. Related to Fig.1a, where a comparable pattern of Endo180 staining in human breast cancers is observed using an independent antibody. **b** Left panel, upregulated expression of genes with *P* value < $10^{-11}$ in CAFs compared to endothelial, immune or epithelial/tumour cell populations from the human colorectal cancer GSE39397 dataset[9]. Right panel, list of the 30 genes commonly upregulated in CAF *vs.* endothelial, CAF *vs.* immune and CAF *vs.* epithelial/tumour cells, with Endo180 (*MRC2*) highlighted. Related to Fig. 1b. **c** tSNE visualisation of stromal cells and colour coded cell types isolated from primary B16-F10 mouse melanoma tumours and draining lymph nodes and subject to single cell RNA-seq (bioRxiv doi.org/10.1101/467225). Expression of marker genes for each cell type, *Mrc2* (bottom left) and *Acta2* (bottom right) analysed from deposited data (https://www.ebi.ac.uk/gxa/sc/experiments/E-EHCA-2/Results). **d** Correlation of Endo180 (*MRC2)* gene expression with a fibroblast TGFβ response signature (F-TBRS)[9] in the luminal A, luminal B, HER2-enriched and basal-like breast cancer intrinsic subtypes in the TCGA dataset. Related to Fig. 1g.

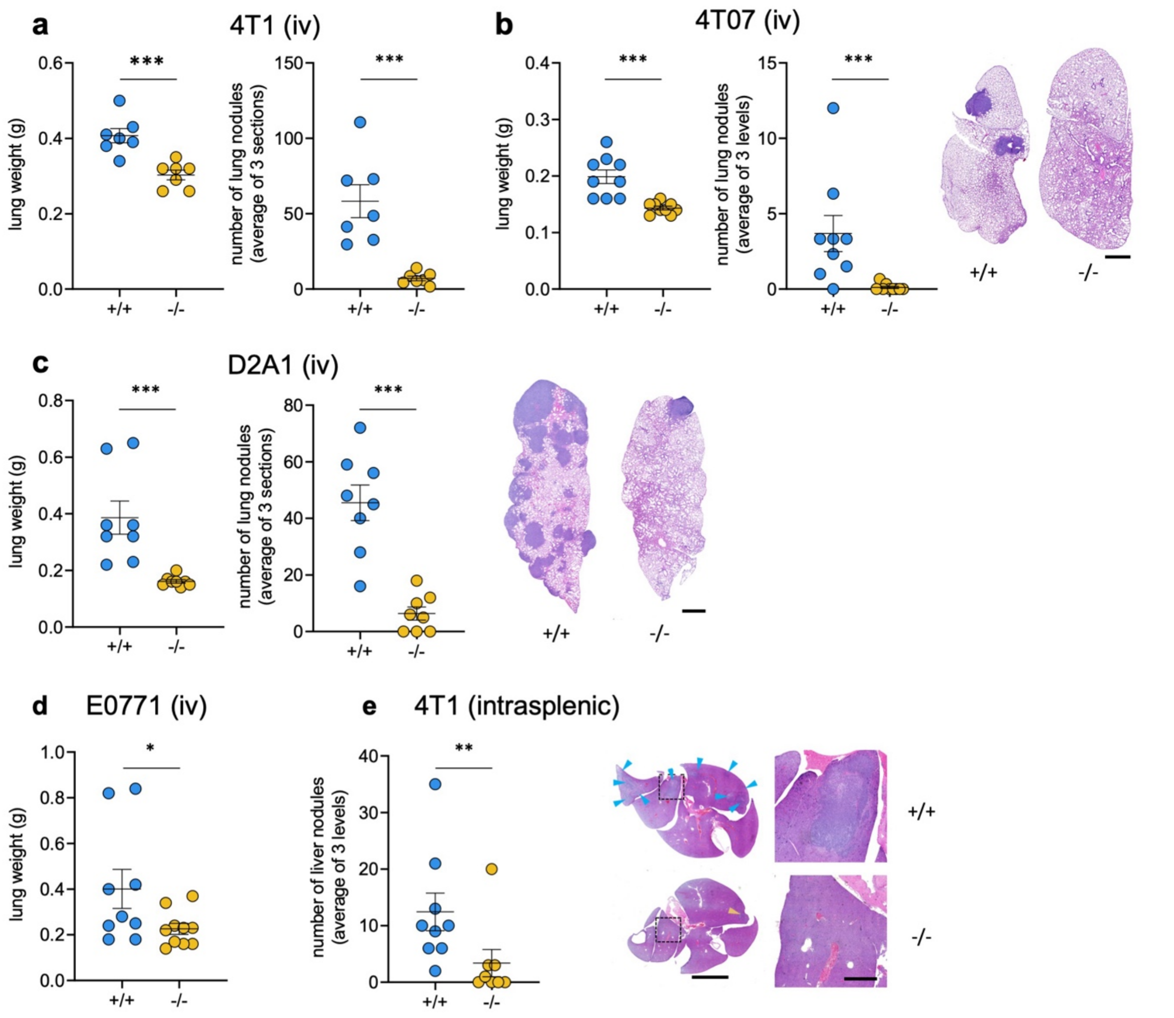


**Supplementary Fig. 2 | Endo180 promotes metastatic tumour growth in lungs and liver**

Additional quantification of the data presented in Figure 3. All data are mean values ±SEM. Quantification of metastatic nodules represents mean number of lung nodules in 3 lung or liver sections. **a** 4T1-Luc cells injected intravenously into BALB/c mice (n = 7 per group) from Figure 3a. Shown are *ex vivo* lung weights (*t*-test) and number of metastatic lung nodules (*t*-test). **b** 4T07 cells injected intravenously into BALB/c mice (n = 9 per group) from Figure 3b. Shown are *ex vivo* lung weights (*t*-test) and number of metastatic lung nodules (Mann-Whitney *U* test, scale bar, 1 mm). **c** D2A1 cells injected intravenously into BALB/c mice (n = 8 per group) from Figure 3c. Shown are *ex vivo* lung weights (Mann-Whitney *U* test), number of metastatic lung nodules (*t*-test) and representative H&E stained sections (scale bar, 1 mm). **d** E0771-Luc cells injected intravenously into C57BL/6 mice (n = 9 or 10 per group) from Figure 3e. Shown are *ex vivo* lung weights (Mann-Whitney *U* test). **e** 4T1-Luc cells injected into the spleen parenchyma of BALB/c mice (n = 8 or 9 per group) from Figure 3f. Shown are number of metastatic liver nodules (Mann-Whitney *U* test) and low (scale bar, 5 mm) and high power (scale bar, 1 mm) images of H&E stained liver sections. Arrowheads indicate tumour nodules.

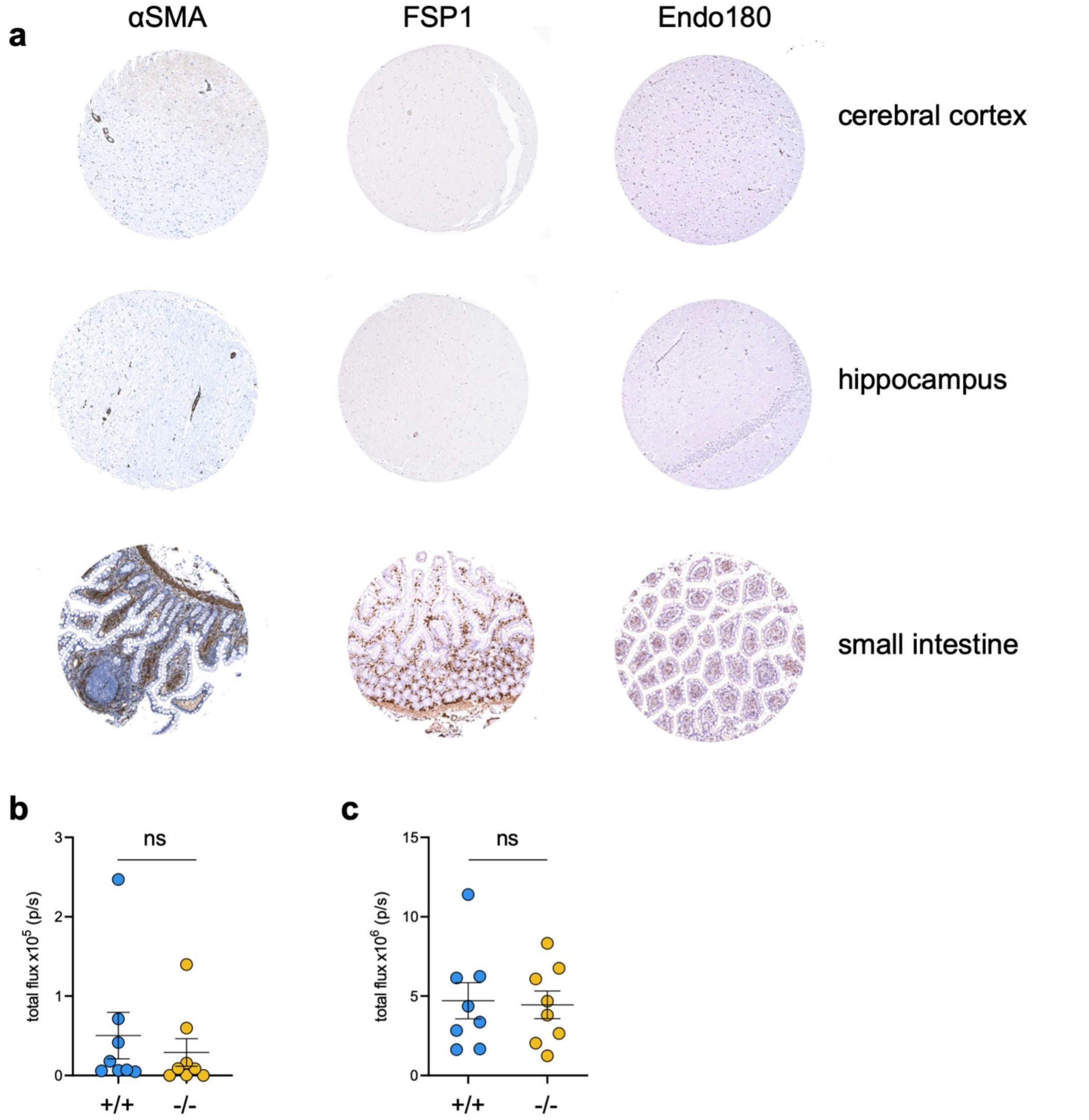


**Supplementary Fig. 3 | Expression of Endo180 in the brain**

**a** Images taken from The Human Protein Atlas of brain (cerebral cortex and hippocampus) and small intestine stained for Endo180 and the fibroblast markers αSMA and FSP1. **b** 2.5 x $10^3$ 4T1-Luc cells were injected intracranially (supraventorial) into BALB/c mice (n = 8 per group). Metastatic colonisation was measured by *ex vivo* IVIS imaging on day 12 (mean values ±SEM, *t*-test, no significant differences). **c** 3 x $10^5$ 4T1-Luc cells were injected into the left ventricle of the heart (n = 8 per group) and metastasis to the brain monitored by *in vivo* IVIS imaging on day 10 (mean values ±SEM, *t*-test, no significant differences). Related to Fig. 3.

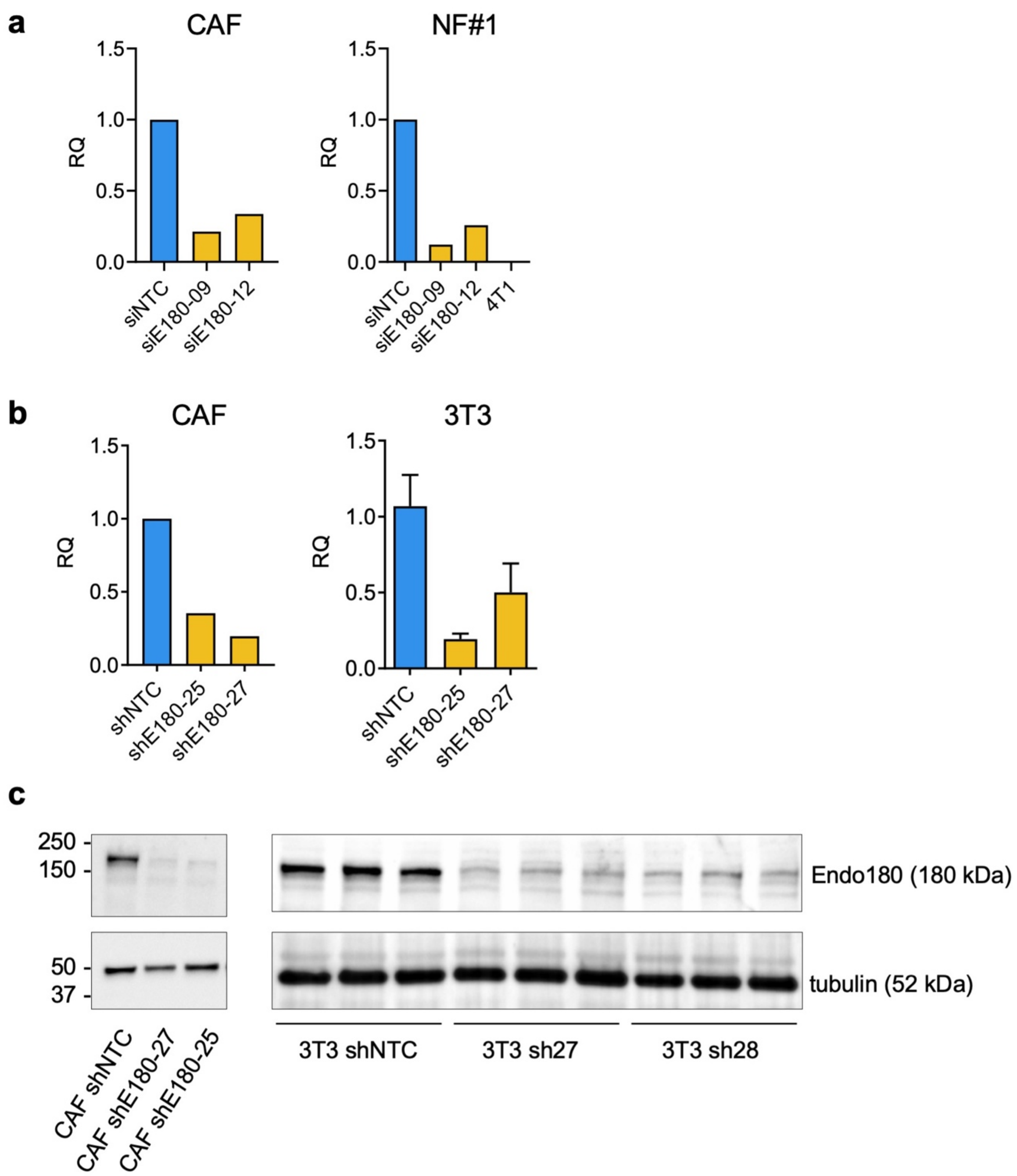


**Supplementary Fig. 4 | Endo180 expression following siRNA- or shRNA-mediated knockdown**

**a, b** Relative Endo180 (*Mrc2*) expression in CAFs and NF#1 and 3T3 mouse fibroblasts after transfection/ transduction with two independent (a) siRNAs or (b) shRNAs in comparison to non-targeting control (NTC) analysed by RT-qPCR. No expression was detected in 4T1 tumour cells. **c** Western blot of Endo180 (180 kDa) in CAFs (left panel) and 3T3 fibroblasts (right panel) transduced with either non-targeting control shRNA (shNTC) or two independent shRNAs against Endo180. Tubulin serves as loading control. Molecular size markers are in kDa. Related to Fig. 4.

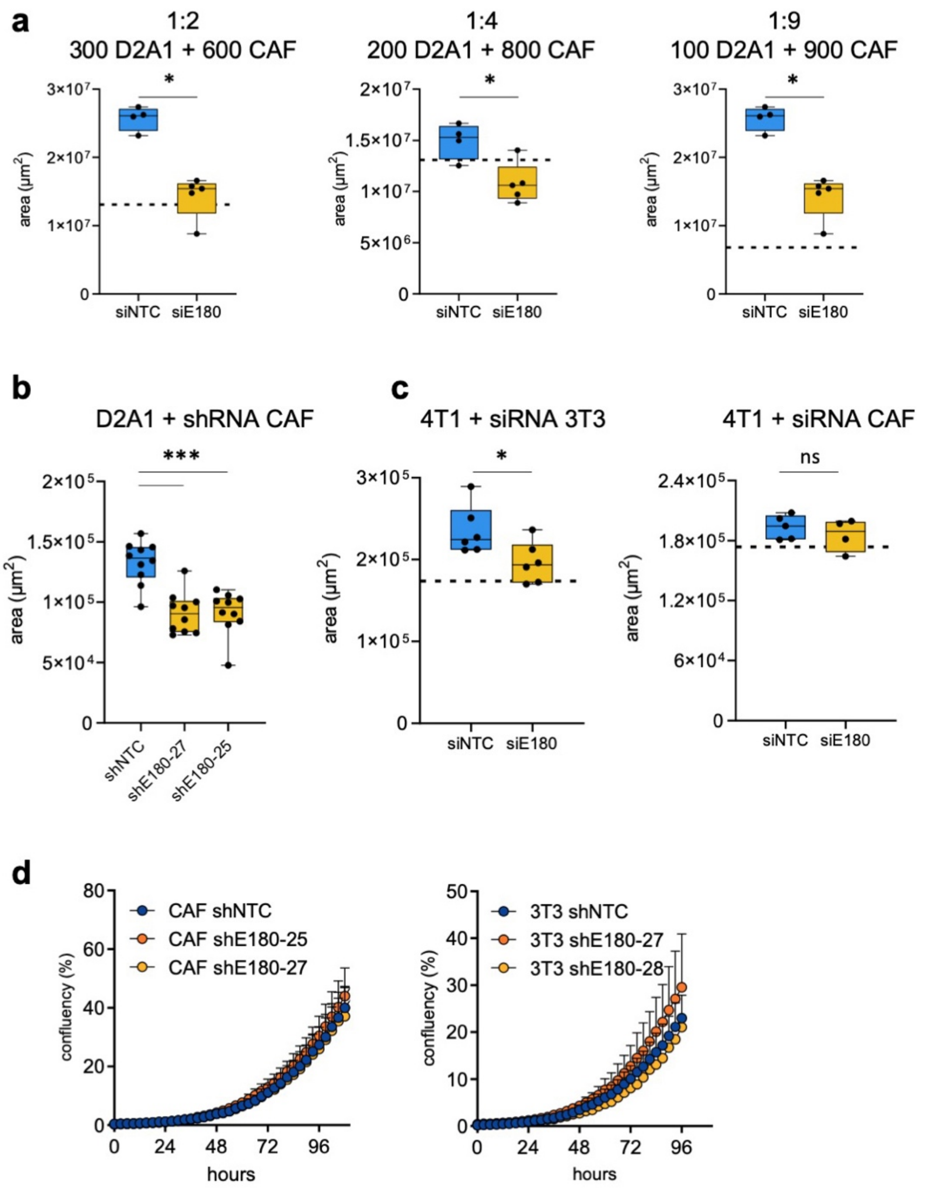


**Supplementary Fig. 5 | Fibroblast-tumour cell co-culture assays**

In all experiments, fibroblasts were transfected with non-targeting control or Endo180 si/shRNAs. **a** Different ratio of D2A1 and CAF cells were co-cultured in U-bottom low adherence plates (n = 4-5 spheroids per condition). Box plot shows median and 25$^{th}$ to 75$^{th}$ quartile, whiskers show minimum and maximum. Mean size of D2A1 spheroids alone is indicated by the dotted line (1:3 ratio, Mann-Whitney *U* test; 1:5 ratio *t*-test, 1:9 ratio Mann-Whitney *U* test). Related to Fig. 4a. **b** Equivalent results to panel a were obtained using CAFs transduced with shRNAs instead of siRNA. Shown are 1:3 ratio co-cultures of D2A1 and CAFs transduced with non-targeting (NTC) or two independent Endo180-targeting shRNAs after 6 days (n = 10 spheroids per condition; mean values ±SEM, one-way ANOVA). Related to Fig. 4a. **c** 4T1 cells admixed with 3T3 or CAFs in U-bottom low adherence plates and cultured for 8 days. Spheroid growth was assessed by spheroid area (n = 4-6 spheres per condition, mean values ±SEM, *t*-test). Related to Fig. 4b. **d** CAFs or 3T3 fibroblasts were cultured in 24-well plates in D2A1 conditioned medium prior to decelluarisation. D2A1 cells were plated onto the fibroblast-derived matrices and growth monitored by IncuCyte imaging (n = 3 wells per condition; mean values ±SEM, 2-way ANOVA). Related to Fig. 4g.

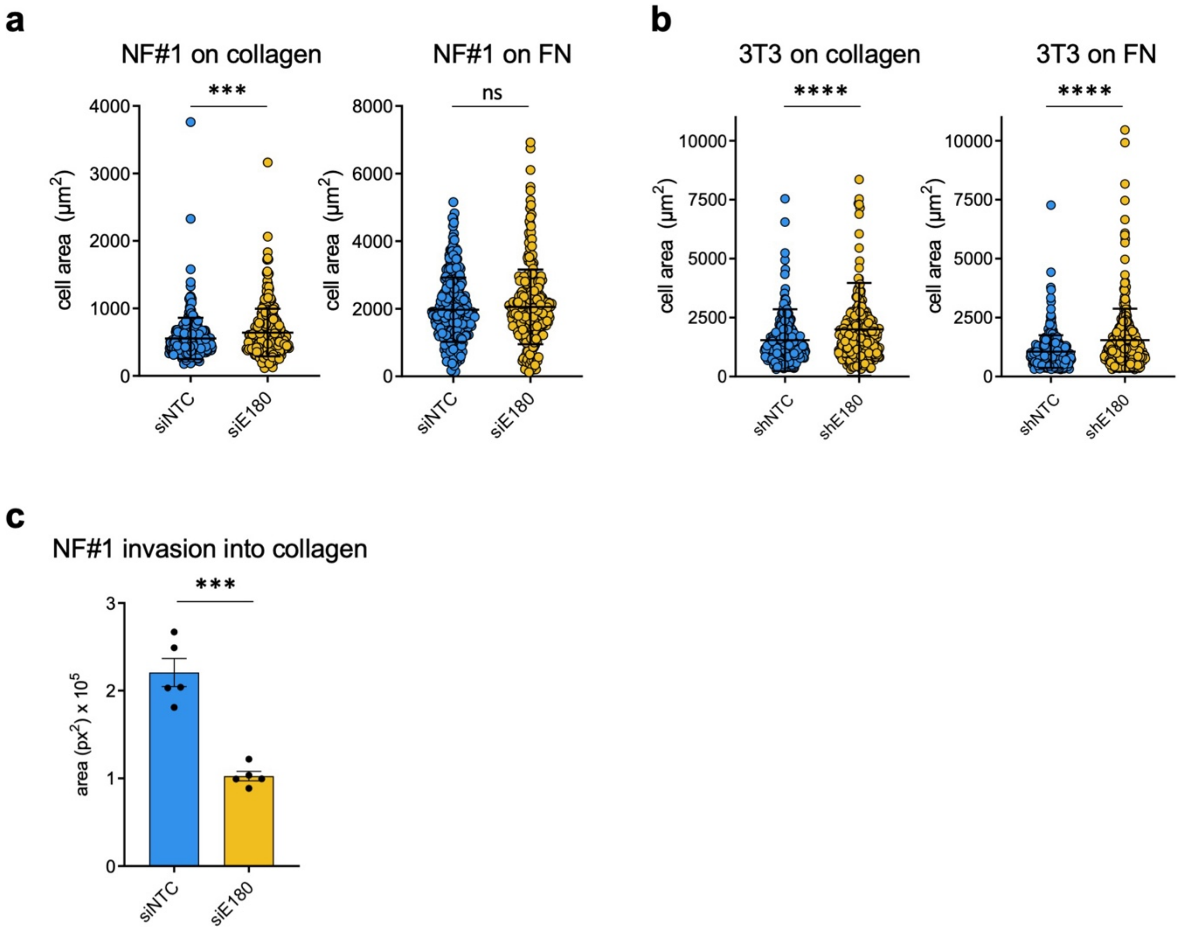


**Supplementary Fig. 6 | Cell spreading on collagen or fibronectin-coated hydrogels**
**a,b** Cell spreading (area) of siNTC and siEnd180 (siE180) NF#1 fibroblasts on collagen or fibronectin-coated hydrogels (50 kPa) and shNTC and shE180-27 3T3 fibroblasts on collagen or fibronectin-coated glass coverslips, respectively, (n = 300 cells; mean values ±SEM, Mann-Whitney *U* test). Related to Fig 5a,b. **c** Invasion of siNTC or siE180 NF#1 fibroblasts from 3D aggregates into a collagen matrix after 48 hours (n = 4; mean values ±SEM, *t*-test). Related to Fig. 5d.

### Supplementary Figure 7

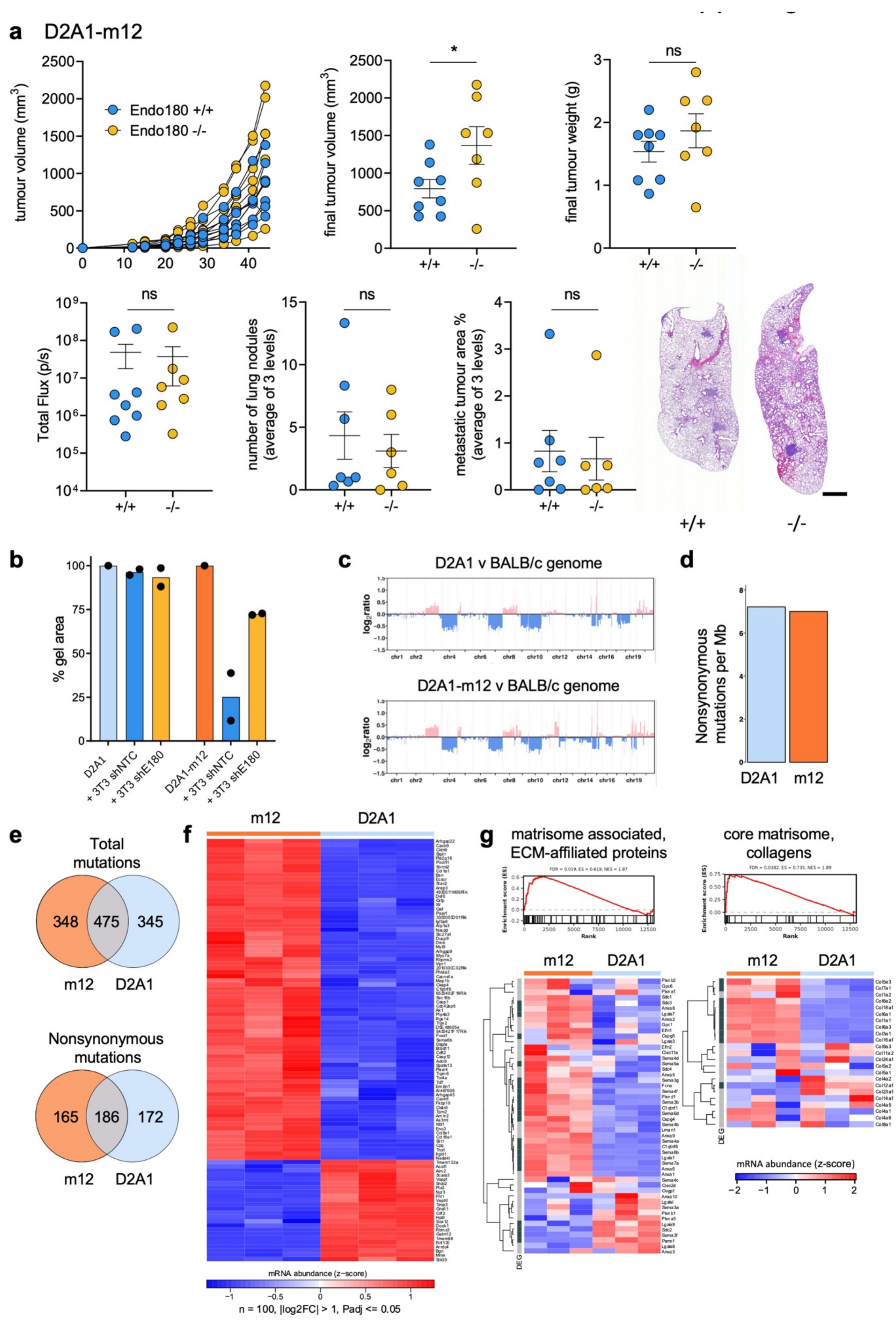

**Supplementary Fig. 7 | Comparison of the parental D2A1 line and the D2A1-m12 subline**

**a** 5 x $10^4$ D2A1-m12-Luc cells were injected into the $4^{th}$ mammary fat pad of BALB/c Endo180$^{+/+}$ or Endo180$^{-/-}$ mice (n = 8 per group) and tumour volume measured twice weekly. The experiment was terminated on day 41. Shown are: D2A1-m12 primary tumour growth in individual mice, final tumour volume, primary tumour weights, quantification of metastatic burden by *ex vivo* IVIS imaging of the lungs, number of lung nodules and metastatic tumour area in the lungs. All data are mean values ±SEM (*t*-test). Representative H&E stained lung sections are shown (scale bar, 1 mm). Related to Fig. 7b. **b** shNTC or shEndo180 3T3 fibroblasts were embedded in a 1:1 ratio (each 5 x $10^5$ mL$^{-1}$) with either D2A1 or D2A1-m12 tumour cells in collagen gels. Data show % gel contraction after 48 hours (n = 2). **c** Copy number variation plots ($\log_2$ ratio) for D2A1 and D2A1-m12 cell lines using a reference BALB/c genome. **d** Tumour mutational burden of D2A1 and D2A1-m12 cell lines as determined using whole-exome sequencing (WES). **d** number of protein-coding nonsynonymous mutations per Mb of exome. **e** Venn diagrams illustrating the number of total mutations and nonsynonymous mutations in common between the D2A1 and D2A1-m12 cell lines. **f** Heatmap displaying mRNA abundance of top 100 significantly differentially expressed genes (DEGs) determined by RNA-seq, in D2A1-m12 compared with D2A1cells (n = 3; heatmap scale is a z-score). Threshold for differential expression was $|\log_2 FC| > 1$, and adjusted *P* value ≤ 0.05. **g** fGSEA of 'core matrisome, collagens' and 'matrisome-associated, ECM-affiliated proteins', and associated heatmaps showing the genes of the pathway (n = 3; heatmap scale is a z-score). Dark grey, significant DEGs with $|\log_2 FC| > 1$ and adjusted *P* value ≤ 0.05. Light grey, nonsignificant DEGs. Related to Fig. 7f.

### References


1. Di Tommaso, P. *et al.* Nextflow enables reproducible computational workflows. *Nat Biotechnol* **35**, 316-319 (2017).
2. Li, H. & Durbin, R. Fast and accurate long-read alignment with Burrows-Wheeler transform. *Bioinformatics* **26**, 589-595 (2010).
3. Li, H. *et al.* The Sequence Alignment/Map format and SAMtools. *Bioinformatics* **25**, 2078-2079 (2009).
4. McKenna, A. *et al.* The Genome Analysis Toolkit: a MapReduce framework for analyzing next-generation DNA sequencing data. *Genome Res* **20**, 1297-1303 (2010).
5. Talevich, E., Shain, A.H., Botton, T. & Bastian, B.C. CNVkit: Genome-Wide Copy Number Detection and Visualization from Targeted DNA Sequencing. *PLoS Comput Biol* **12**, e1004873 (2016).
6. Wilm, A. *et al.* LoFreq: a sequence-quality aware, ultra-sensitive variant caller for uncovering cell-population heterogeneity from high-throughput sequencing datasets. *Nucleic Acids Res* **40**, 11189-11201 (2012).
7. Wang, K., Li, M. & Hakonarson, H. ANNOVAR: functional annotation of genetic variants from high-throughput sequencing data. *Nucleic Acids Res* **38**, e164 (2010).
8. Wienke, D. *et al.* The collagen receptor Endo180 (CD280) Is expressed on basal-like breast tumor cells and promotes tumor growth in vivo. *Cancer Research* **67**, 10230-10240 (2007).
9. Calon, A. *et al.* Dependency of colorectal cancer on a TGF-beta-driven program in stromal cells for metastasis initiation. *Cancer cell* **22**, 571-584 (2012).
